# Molecular determinants of MED1 interaction with the DNA bound VDR-RXR heterodimer

**DOI:** 10.1101/2020.06.17.157305

**Authors:** Anna Y. Belorusova, Maxime Bourguet, Steve Hessmann, Sandra Chalhoub, Bruno Kieffer, Sarah Cianférani, Natacha Rochel

**Author notes:** Co-first authors (Anna Y. Belorusova, Maxime Bourguet). Correspondence to Anna Y. Belorusova or Natacha Rochel.

## Abstract

The MED1 subunit of the Mediator complex is an essential coactivator of nuclear receptor-mediated transcriptional activation. While structural requirements for ligand-dependent binding of classical coactivator motifs of MED1 to numerous nuclear receptor ligand-binding domains have been fully elucidated, the recognition of the full-length or truncated coactivator by full nuclear receptor complexes remain unknown. Here we present structural details of the interaction between a large part of MED1 comprising its structured N-terminal and the flexible receptor-interacting domains and the mutual heterodimer of the vitamin D receptor (VDR) and the retinoid X receptor (RXR) bound to their cognate DNA response element. Using a combination of structural and biophysical methods we show that the ligand-dependent interaction between VDR and the second coactivator motif of MED1 is crucial for complex formation and we identify additional, previously unseen, interaction details. In particular, we identified RXR regions involved in the interaction with structured N-terminal domain of MED1, as well as VDR regions outside the classical coactivator binding cleft affected by coactivator recruitment. These findings highlight important roles of each receptor within the heterodimer in selective recognition of MED1 and contribute to our understanding of the nuclear receptor-coregulator complexes.

## Introduction

Nuclear receptor (NR) superfamily of ligand-regulated transcription factors activate or repress gene expression by recruiting coactivators, corepressors, chromatin remodelers and the general transcriptional machinery to the target genes (reviewed in Lonard & O’Malley, 2012 and Giudici et al., 2015). Among classical NR coactivators is Mediator, an evolutionary conserved multi-protein complex facilitating multiple stages of gene expression, notably the chromatin remodeling and pre-initiation complex formation (reviewed in Jeronino and Robert 2017; Soutourina 2018; Verger et al. 2019). It was discovered as a group of factors needed for the yeast RNA Polymerase II activity (Kelleher et al., 1990; Flanagan et al., 1991), and subsequently various mammalian Mediator subcomplexes have been isolated through association with NRs, such as TRAP complex associated with thyroid receptor (TR), (Fondell et al., 1996) and DRIP complex associated with vitamin D receptor (VDR) (Rachez et al., 1998). Other similar complexes included activator-recruited cofactor ARC (Näär et al., 1999), mammalian mediator (Jiang et al., 1998), mammalian Srb/Mediator complex (Boyer et al., 1999), PC2 (Malik et al., 2000) and CRSP (Ryu and Tjian 1999). Mediator is involved in strong ligand-dependent interaction with NRs primarily via its largest subunit 1 (MED1) (Yuan et al. 1998; Rachez et al. 1999; Zhu et al. 1997; Kang et al. 2002; Ge et al. 2002; Hittelman et al. 1999; Malik et al. 2002; Wang et al. 2002), although for some NRs the interaction can include other Mediator subunits and alternative cofactors (Kim et al. 2008; Chen et al. 2009; Wallberg et al. 2003; Iida et al. 2015; Harms et al. 2015).

As a classical NR-binding target, MED1 contains two LXXLL motifs, also called NR-boxes, localized in a central disordered receptor-interacting domain (RID). Binding of the coactivator LXXLL motifs to the activation function 2 (AF-2) of the receptor ligand-binding domain (LBD) has been extensively characterized by structural studies (Nolte et al., 1998; Shiau et al., 1998; Darimont et al., 1998). Leucines from the coactivator LXXLL motif are buried in the hydrophobic groove of the AF-2 surface formed by hydrophobic residues from helices H3, H4 and H12 of the LBD, and the NR box is locked by a charge clamp formed by a lysine on the NR H3 and a glutamate on H12. The two MED1 LXXLL motifs bind to NRs with different specificity: steroid hormone receptors preferentially bind to the first LXXLL motif, whereas non-steroid hormone receptors, such as TR and VDR, strongly interact with the second LXXLL motif (Burakov et al., 2000; Yuan et al., 1998; Ren et al., 2000).

Mediator-dependent mechanisms of NR regulation by MED1 include looping of enhancers to transcription start sites via an assembly process involving transcription factors, cohesin and non-coding RNAs (Lai et al. 2013; Hsieh et al. 2014; Step et al. 2014; Kagey et al. 2010 ; Sanyal et al. 2012); directly linking chromatin remodeling and the pre-initiation complex formation (Chen et al., 2009; Wallberg et al., 2009); or repression of transcription through the core Mediator-associated CDK8 kinase module (reviewed in Chen & Roeder, 2011). At the same time, some regulatory roles of MED1 could be Mediator-independent as it can be recruited to NRs independently from the other Mediator subunits, promoting the association with the Mediator core in a second step. Occupancies of MED1 and NRs on the genome sites are highly correlated (Meyer and Pike, 2013; Step et al., 2014) and MED1 levels are highly elevated on super-enhancers in embryonic stem cells and in cancer cells where NRs act as master regulators (Whyte et al., 2013; Lovén et al., 2013). Recent studies showed that intrinsically disordered regions of MED1 can form phase-separated droplets that compartmentalize and concentrate transcriptional regulators (Sabari et al., 2018).

MED1 has been shown to be essential for various biological functions of large number of NRs (reviewed in Chen & Roeder, 2011). Due to its important role in human physiology, it has been suggested as a possible target for several disorders, such as metabolic syndrome (Chen et al. 2010), fatty liver (Bai et al., 2011) and several types of cancer, including breast and prostate cancers (reviewed in Weber & Garabedian, 2017).

Despite essential role and high therapeutic potential of MED1, no atomic structural data is available for this protein or its homologs. Furthermore, while much has been discovered about the Mediator complex and its association with transcriptional machinery, the mechanistic details of how MED1 bridges the Polymerase II to NRs are far less understood. Most of the structural investigations on NR-MED1 association, similarly to analogous coregulator complexes, are still limited to the recognition of LXXLL-peptides or short RIDs by NRs (Osz et al., 2012; Rochel et al., 2011; Pavlin et al., 2014). Recent advances in single particle cryo-electron microscopy (cryoEM) allowed the structural characterization of the full-length ERα−p160/p300 coactivator complexes (Yi et al., 2015; Yi et al., 2017). However, the detailed mechanism of how NRs trigger the formation of big regulatory complexes that directly alter the transcriptional rate has not yet been fully elucidated and remains challenging due to the presence of large intrinsically disordered regions in the coactivators proteins and the associated flexibility of the complexes.

To provide structural insights into the mechanism of the NR-MED1 specific association, in the present study we investigated the complex formed between a large fragment of the coactivator MED1 comprising its structured N-terminal region and the RID encompassing two LXXLL motifs and the full NR heterodimer formed by VDR and the Retinoid X Receptor (RXR). We combined structural methods including small angle X-ray scattering (SAXS), NMR, hydrogen-deuterium exchange coupled to mass spectrometry (HDX-MS), crosslinking mass spectrometry (XL-MS) as well as biophysical methods to characterize the MED1 recruitment by the receptor heterodimer and to get structural details of this assembly. We show that one molecule of MED1 is recruited by the VDR-RXR heterodimer and confirm primary role of the VDR AF-2 interaction with the second LXXLL motif of MED1 in complex formation. We demonstrate that the RXR AF-2 is not essential for the MED1 recruitment, however is affected upon MED1 binding. We also identify other RXR regions, as well as VDR regions outside the AF-2, which are included in the interaction and could be important for reaching the coactivator selectivity by VDR-RXR. Novel structural information on the NR-MED1 complex presented in this work is essential to understand the molecular organization and the interaction networks between complexes of such type.

## Material and Methods

### Compounds

1α,25-dihydroxyvitaminD3 (1,25D3) and 9cis retinoic acid (9cisRA) were purchased from Sigma. The *rANF* single strands DNAs (5’-AGAGGTCATGAAGGACATT-3’ and 5’AATGTCCTTCATGACCTCT-3’) were purchased from Sigma Aldrich and annealed. The MED1 NR2 peptide (NHPMLMNLLKDN) was synthesized by Pascal Eberling (IGBMC peptide synthesis common facility).

### Biochemistry

In all experiments, HsVDR-HsRXRA complexes were analyzed, except for surface plasmon resonance experiment where HsVDR-*Mm*RXRA complexes were used. cDNAs encoding full-length hVDR (1-427), hVDRΔ166-216(1-427, Δ166-216) and hVDRΔH12 (1-415) cloned into the pET28b vector were used to generate the N-terminal His-tagged proteins. hRXRαΔNTD (130-462), mRXRαΔNTD (132-467), mRXRαΔNTDΔH12 (132-449), mRXRαΔNTD K289A,E458A cloned into pET15b, were expressed as N-terminal His-tagged proteins. Recombinant proteins were produced in Escherichia coli BL21 DE3 after induction with 1 mM IPTG (OD600 ∼ 0.7) at 23 °C for 4 hours. Soluble proteins were purified using chromatography column (HisTrap FF crude, 17-5286-01, GE) followed by size exclusion chromatography (SEC) on HiLoad Superdex 200 (28-9893-35 GE) equilibrated in 20 mM Tris-HCl, pH 8.0, 250 mM NaCl, 5% glycerol, 2 mM CHAPS and 1 mM TCEP. Full-length hVDR and hRXRαΔNTD were mixed in stoichiometric amounts and purified by size exclusion chromatography (HiLoad Superdex 200, 28-9893-35, GE) equilibrated in 20 mM Tris-HCl, pH 8.0, 200 mM NaCl, 5% glycerol, 2 mM CHAPS and 1 mM TCEP. Ligands (1,25D3 and 9cisRA) were added to the stoichiometric heterodimer and *rANF* DR3 was mixed in a 1.1 equivalent ratio. The DNA complex was further purified by Size Exclusion Chromatography (SEC) in 20 mM Tris pH 8.0, 75 mM NaCl, 75 mM KCl, 2 mM CHAPS, 5% Glycerol, 4 mM MgSO_4_, 1 mM TCEP. A cDNA encoding truncated hMED1 (50-660) cloned into pBacHGW, pFastBac-1 (InVitrogen) baculovirus transfer vector adapted for Gateway, was used to produce MED1 proteins with N-terminal His-tag. Sf9 cells were infected with recombinant baculovirus at a multiplicity of infection equal 5 and cultured in TNM-FH supplemented with 10% FCS and 50 mg/ml gentamycin at 27°C for 48 h. Cells were harvested by centrifugation (1,000 g for 15 min) and cell pellets were stored at -20°C prior purification. Soluble protein was purified using batch/gravity-flow affinity chromatography (cOmplete, Roche). MED1 (50-660) was eluted by 250 mM Imidazole in binding buffer. Following the His-tag removal by Thrombin cleavage, the protein was further purified by SEC on HiLoad Superdex 200 (28-9893-35 GE) equilibrated in 20 mM Tris-HCl, pH 8.0, 250 mM NaCl, 5% glycerol, 2 mM CHAPS and 1 mM TCEP.

The proteins were concentrated to 3-6 mg/ml with an Amicon Ultra 30 kDa MWCO. Purity and homogeneity of all proteins were assessed by SDS and Native PAGE.

### Gel retardation in TBE

6% polyacrylamide gel was used to examine the migration of DNA-bound complexes and MED1. The samples were loaded onto the polyacrylamide gel, placed in a Bio-Rad chamber for gels and ran with constant current of 6 mA for 3 hours at 4°C in TBE migration buffer. Gels were revealed by Coomassie staining.

### Small angle X-ray scattering

Synchrotron X-ray data were collected on a Pilatus 1M detector at the ESRF beamline BM29 (Pernot et al., 2010). 100 μL of VDR-RXRΔNTD-DR3, MED1 (50-660) and their complex at concentrations 8.5 - 10 mg mL^-1^ in 25 mM HEPES pH 7.5, 150 mM NaCl, 5% Glycerol, 2 mM MgCl2, 2 mM TCEP onto a GE Healthcare Superdex 200 10/300 column (equilibrated in the same buffer) at a flow rate of 1 mL.min^-1^. A scattering profile was integrated every second. Frames were selected based on the examination of the SEC profile together with the calculated Rg and Dmax values. The SAXS data were averaged and processed by standard procedures using PRIMUS (Konarev et al., 2003). The forward scattering I(0) and the radii of gyration Rg were evaluated using the Guinier approximation assuming that at very small angles (s < 1.3/Rg) the intensity is represented as I(s) = I(0) exp(-(sRg)^2^/3). These parameters were also computed from the entire scattering pattern using the indirect transform package GNOM (Svergun, 1992) which also provides the maximum dimension of the particle Dmax and the distance distribution function P(r). The program SASREF (Petoukhov and Svergun, 2005) was employed for molecular rigid body modeling of the RXR/VDR/DNA complex, based on SAXS and cryoEM structures (Rochel et al., 2011; Orlov et al., 2012). The final fits of the model scattering to the experimental data were computed using CRYSOL (Svergun et al., 1995).

### SEC-MALLS

The molecular weight and homogeneity of the sample was checked using a SEC column coupled with MALLS Dawn DSP detector (Wyatt Technology, Santa Barbara, CA, USA). A GE Healthcare Superdex 200 10/300 analytical column was pre-equilibrated with the sample buffer, 25 mM HEPES pH 7.5, 150 mM NaCl, 5% Glycerol, 2 mM MgCl2, 2 mM TCEP. The system was operated at 20°C, with a flow rate of 0.75 ml/min.

### Analytical ultracentrifugation

Sedimentation velocity experiments were performed at 4°C in 25 mM Tris-HCl pH 8.0, 100 mM NaCl, 2% Glycerol, 1 mM CHAPS, 2 mM MgCl2, 1 mM TCEP using Beckman Coulter Proteome Lab XL-I analytical ultracentrifuge and the 8-hole Beckman An-50Ti rotor. Sedimentation at 50,000 rpm was monitored by absorbance at 280 nm with boundaries measured each 7 min. MED1 (50-660) at constant concentration (6 μM) was titrated by VDR-RXR-*rANF1*; tested ratios varied from 1:0.5 to 1:2.7. Density and viscosity of the used buffer were calculated using SEDNTERP software (http://sednterp.unh.edu/) and used for the data correction. Using nonlinear least-squares analysis with SEDPHAT (Schuck, 2000), collected datasets were fitted using single site heter-association model.

### Hydrogen Deuterium Exchange (HDX) coupled to mass spectrometry (MS) experiments

HDX experiments of VDR-RXRΔNTD-*rANF1* complex were carried out with and without 2 molar excess of NR2 motif in 20mM Tris pH 8.0, 75mM NaCl, 75mM KCl, 2mM CHAPS, 5% glycerol, 4mM MgSO4, 1mM TCEP. The same buffer was used for VDR-RXR-*rANF1*-MED1(50-660) complex HDX experiment. Preparation and injection of the samples were automatically conducted using a LEAP HDX Automation Manager (Waters), while chromatography was carried out on an Acquity UPLC system with HDX technology (Waters, Manchester, UK). Samples were incubated at different deuteration times (0⍰, 0.5⍰, 2⍰, 10⍰ and 30⍰min) in 95% of deuterated buffer (20⍰mM Tris pD 8.0, 75 mM NaCL, 75 mM KCl, 4 mM MgSO4, 1 mM TCEP) before quenching the exchange by adding a 150⍰mM glycine pH 2.4, 2⍰M GdHCl, 4mM MgSO4, 1mM TCEP buffer at 1⍰°C during 30⍰s. Digestion of samples (between 20 and 50 pmol injections) was then performed through a pepsin-immobilized cartridge in 0.1% aqueous formic acid solution at a 200⍰µl/min. Generated Peptides were then trapped on a UPLC pre-column (ACQUITY UPLC BEH C18 VanGuard pre-column, 2.1⍰mm I.D. ⍰×⍰5⍰mm, 1.7⍰µM particle diameter, Waters) and separated on UPLC column (ACQUITY UPLC BEH C18, 1.0⍰mm I.D. ⍰×⍰100⍰mm, 1.7⍰µM particle diameter, Waters) at 0⍰°C. Mass spectrometry analyses were acquired with Synapt G2Si HDMS (Waters) with electrospray ionization, using data-independent acquisition mode (MS^E^) over an *m*/*z* range of 50–2000 and 1001fmol/µl Glu-FibrinoPeptide solution as lock-mass correction and calibration. Analyses were performed with the following parameters: capillary voltage, 3 kV; sampling cone voltage, 40 V; source temperature, 80°C; desolvation gas, 150°C and 600 L.h^-1^; scan time, 0.3 s; trap collision energy ramp, 15 to 40 eV. HDX experiments were realized in triplicate for each time point. Peptide identification was performed using ProteinLynx Global Server 2.5.3 (Waters) with a home-made protein sequence library containing VDR, RXR and MED1(50-660) sequences, with peptide and fragment tolerances set automatically by PLGS, and oxidized methionine set as variable modification. Deuterium uptakes for all identified peptides were then filtered and validated manually using DynamX 3.0 (Waters) as follows: only peptides identified in all replicates were kept with only one charge state with a minimum fragment of 0.2 per amino acid, a minimum intensity at 103, a length between 5 and 30 residues and a file threshold of 3. Deuterium uptakes were not corrected and are reported as relative. HDX–MS results were statistically validated using Mixed-Effects Model for HDX experiments (MEMHDX, Hourdel et al., 2016) where statistical significance thresholds were set to 0.01. HDX results were exported on VDR-RXR SAXS model using PyMOL (www.pymol.org). HDX-MS data have been deposited to the ProteomeXchange Consortium via the PRIDE (Perez-Riverol et al., 2019) partner repository with the dataset identifier PXD019530.

### Chemical crosslinking (XL) coupled to mass spectrometry (MS) experiments

Crosslinking reactions were conducted with 25 µM protein solutions in 20 mM HEPES pH 8.0, 75 mM NaCl, 75 mM KCl, 4 mM MgSO_4_, 1 mM TCEP Freshly prepared 10 mM stock solution of DSBU and C2-arm version of DSBU in DMSO (CF Plus Chemicals s.r.o., Czech Republic) were added in 50-, 100- and 200-fold molar excess. Crosslinking reactions were conducted during 45 min at room temperature and furtherquenched during 20 min using NH_4_HCO_3_ to a final concentration of 20 mM final. Disulfide reduction was next performed by incubating the crosslinked complex solution with 5 mM DTT for 30 min at 60°C, followed by alkylation with 15 mM IAA for 30 min in the dark. Then trypsin (Promega, Madison, USA) was added in 1:50 enzyme:substrate ratio. Samples were incubated overnight at 37°C. Digestion was quenched with 1% formic acid. Peptides were cleaned up using SPE cartridges and samples were concentrated in a SpeedVac concentrator before LC/MS/MS analysis. NanoLC-MS/MS analyses were performed using a nanoAcquity UPLC (Waters, Milford, USA) coupled to the Q-Exactive Plus Orbitrap mass spectrometer (Thermo Scientific, Bremen, Germany) Nanospray Flex™ Ion source. The samples were trapped on a nanoACQUITY UPLC precolumn (C18, 180µm x 20mm, 5 µm particle size), and the peptides were separated on a nanoACQUITY UPLC column (C18, 75µm x 250 mm with 1.7 µm particle size, Waters, Milford, USA) maintained at 60°C. The samples were first injected with a 285 min gradient and a flow rate of 450 nL/min. The Q-Exactive Plus Orbitrap source temperature was set to 250°C and spray voltage to 1.8kV. Full scan MS spectra (300-1800 m/z) were acquired in positive mode at a resolution of 140 000, a maximum injection time of 50 ms and an AGC target value of 3 × 10^6^ charges, with lock-mass option being enabled (polysiloxane ion from ambient air at 445.12 m/z). The 10 most intense multiply charged ions per full scan (charge states ⍰2) were isolated using a 2 m/z window and fragmented using higher energy collisional dissociation (30 normalized collision energy, ±3%). MS/MS spectra were acquired with a resolution of 35 000, a maximum injection time of 100 ms,an AGC target value of 1 x 10^5^ and dynamic exclusion was set to 60 sec. The system was fully controlled by XCalibur software v3.0.63, 2013 (Thermo Scientific) and NanoAcquity UPLC console v1.51.3347 (Waters). Raw data collected were processed and converted into .mgf format. The MS/MS data were analyzed using MeroX software version 1.6.6 (Götze et al 2015). Mass tolerence of 5 ppm for precursor ions and 10 ppm for product ions were applied. A 5% FDR cut-off and a signal-to-noise ≥2 were applied. For both crosslinkers, Lys and Arg were considered as protease cleavage sites with a maximum of three missed cleavages. Carbamidomethylation of cysteine was set as fixed and oxidation of methionine as variable modifications (max. mod. 2). Primary amino groups (Lys side chains and N-termini) as well as primary hydroxyl groups (Ser, Thr and Tyr side chains) were considered as crosslinking sites. The cRap database was used in combination with the reporter ion scan event (RISE) mode on. Crosslinks composed of consecutive amino acid sequences were not considered. Each crosslinked product automatically annotated with MeroX was manually validated. Finally, PyMOL software (www.pymol.org) was used to calculate the Cα-Cα distance of each validated linkage sites. The Xl-MS data set has been deposited to the ProteomeXchange Consortium via the PRIDE (Perez-Riverol et al., 2019) partner repository with the dataset identifier PXD019530.

### NMR

NMR experiments were recorded at 280 K on an Avance III Bruker 700 MHz equipped with a z-gradient TCI cryoprobe. NMR samples consisted of 65 µM solution of ^15^N labelled VDR, either full-length or VDRΔ166-216, alone or in complex with 1,25D3, liganded RXRΔNTD and *rANF1* DNA in buffer containing 25 citrate, pH 6.3, 100 mM NaCl, 5% Glycerol, 2 mM MgCl2 and 1 mM TCEP in a 3 mm NMR tube. Interaction with MED1 was studied by adding equimolar amounts of MED1 (50-660) protein to the VDR-RXRΔNTD-DNA and VDRΔ166-216-RXRΔNTD-DNA complexes, where VDR was full-length or devoid of the LBD insertion region, respectively. ^1^H-^15^N HSQC were recorded using WATERGATE solvent suppression pulse sequence from the Bruker standard library with a total acquisition time of four hours.

### Surface Plasmon Resonance

Measurements were performed by Biacore T100 sensitivity enhanced T200 equipment (GE Healthcare) using CM5 series S sensor chip (GE) (29-1496-03). MED1(50-660) was immobilized on the chip surface using standard amino-coupling protocol in 10 mM Na-acetate buffer pH 5.5. The resulting immobilized MED1 was in the range of 100-200 response unit. The running buffer was 50 mM Hepes pH 7.5, 400 mM NaCl, 1 mM TCEP, 0.005% Tween 20 and for regeneration 1M sodium chloride solution was used. Interactions of the MED1 with fully liganded VDR-RXR wild type, VDRΔH12-RXR, VDR-RXR AF2 mutant and VDR-RXR ΔH12 were analyzed in the manner of dose response using twofold dilution series of VDR-RXR ranging from 0.01 to 8µM. The association phase was 120 s and the dissociation phase was 120 s. After subtracting the reference and buffer signal, the data were fit to a steady state binding model using the Biacore T200 Evaluation software (GE Healthcare).

## Results

### Binding of MED1 N-terminal domain to VDR-RXR

Previous structural studies uncovered details of the VDR LBD interaction with the second LXXLL motif of MED1 (NR2 motif, residues 645-649) (Ren et al., 2000; Teichert et al., 2009) (Supplementary Figure 1a), and we have previously shown that the MED1 RID binds to the VDR-RXR heterodimer asymmetrically and remains flexible (Rochel et al., 2011). To gain further insights into specific association between VDR-RXR and MED1, we investigated the binding of the receptor heterodimer to a larger fragment of the coactivator. Based on the disorder prediction for MED1 (Supplementary Figure 1a,b), we selected a protein construct spanning from residues 50 to 660 encompassing the structured N-terminus and RID, as previous studies suggested that first 570 residues of MED1 are sufficient for its incorporation into the Mediator (Ge et al., 2008).

We investigated how purified MED1 (50-660) fragment interacts with the heterodimer formed by the full-length VDR and the RXRα lacking the flexible N-terminal domain (NTD), named hereafter VDR-RXR, and associated with a DR3 DNA response element. To stabilize the VDR-RXR heterodimer, we selected the VDRE from *rANF1* gene. Indeed, we showed that VDR-RXR binds to this DNA element with high affinity (Supplementary Figure 2a) in good agreement with previous characterization of the *rANF1* promoter (Kahlen & Carlberg, 1996). In addition, we showed by a transactivation assay in HEK 293 cells that the luciferase gene under control of the *rANF1* promoter is activated by the VDR natural ligand, 1,25D3, at nanomolar concentrations (Supplementary Figure 2b). The complex was formed by an overnight incubation of the purified VDR-RXR bound with their cognate ligands, 1,25D3 and 9cisRA, and the purified MED1 fragment. The presence of the two ligands in the VDR-RXR complex was confirmed by native ESI-MS (data not shown). The complex formation with MED1 was observed by gel retardation assay (Supplementary Figure 2c) and SEC, where the newly formed high molecular weight complex could immediately be detected as a new specimen appearing on the elution profile before the MED1 (68 kDa) and the VDR-RXR (86 kDa) (Supplementary Figure 2d-e). The molecular weight of the complex as measured by multi-angle light scattering was in close agreement with the respective calculated molecular weight of complex where MED1 binds to VDR-RXR in a 1:1 stoichiometry (Figure 1a). The binding mode of MED1 to VDR-RXR-DNA was further analyzed by analytical ultracentrifugation in sedimentation velocity mode (Supplementary Figure 3). The distribution of sedimentation coefficients *c(s)* (Figure 1b) confirmed the formation of the complex between the MED1 and VDR-RXR-DNA and the complex stoichiometry. Interestingly, the *c(s)* peak shifted in a concentration-dependent way indicating fast kinetics of the interaction (k_off_ > 10^−2^ s^-1^).

**Figure 1.**
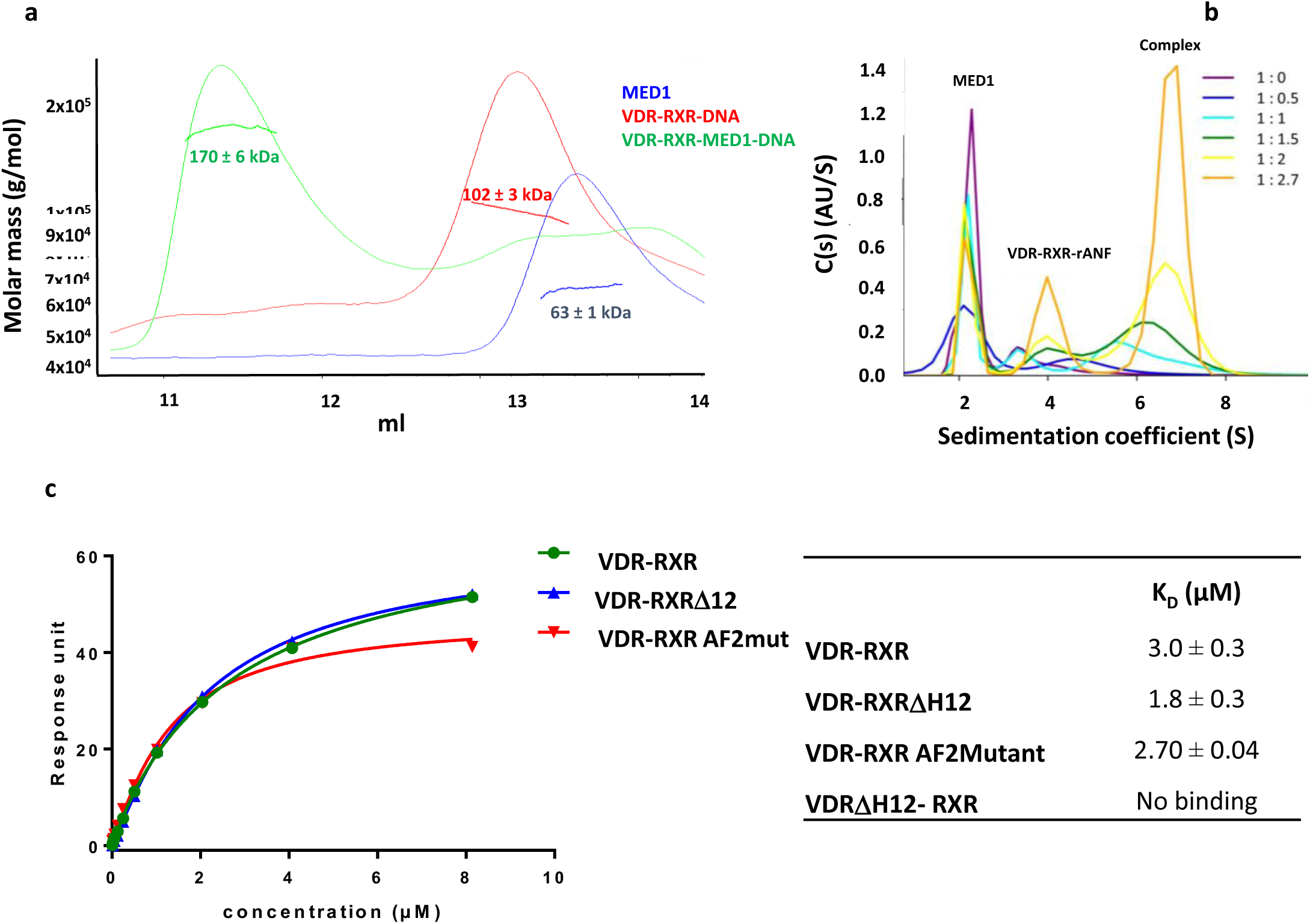
MED1 forms a complex with VDR-RXR-*rANF1* DNA. **(a)** Size-exclusion chromatography-coupled multi-angle laser light scattering (SEC-MALLS) of MED1, VDR-RXR-DNA and VDR-RXR-DNA-MED1 complexes showing the elution profile on a SEC S200 10/300 with the direct molar mass measurement of each elution peak. **(b)** Analytical ultracentifugation: *c(s)* distributions. Ratio MED1:VDR-RXR-DNA is indicated and is used for the color representation of the distributions. Left side of the graph corresponds to the top, and right side – to the bottom of the sample cell, which correlates with the direction of migration. (*c*) Analysis of the interactions of VDR-RXR wild type and VDR-RXRΔH12, VDR-RXR AF2Mutant and VDRΔ12-RXR mutants with MED1 (50-660) in the presence of 1,25D3 and 9-cis RA by surface plasmon resonance and calculated K_D_.

All together, the biophysical data indicates that only one MED1 molecule binds to the liganded VDR-RXR heterodimer and we show that the process of the protein association-dissociation is dynamic and the complex is rather transient.

### Topology of the MED1-VDR-RXR-DNA complex

Small-angle X-ray scattering (SAXS) data were collected from samples of MED1, liganded VDR-RXR-DNA and liganded VDR-RXR-DNA-MED1 using on-line size exclusion chromatography (SEC-SAXS) (Supplementary Figure 4). The SAXS profiles are shown in Figure 2a-b and Supplementary Figure 5 and the structural parameters including the radius of gyration, *R*_*g*_, and the maximum particle dimension, *D*_*max*_, are reported in Table 1. For MED1 alone, the Kratky plot representation of MED1 scattering data (Supplementary Figure 5c) suggests that the fragment is globular and has one core with flexible parts/linkers. The structural parameters values for VDR-RXR-DNA are slightly larger than those previously determined for the related VDRΔ-RXR-DNA complex where VDR-specific insertion localized between H1 and H3 was truncated (Rochel et al., 2011). Binding of MED1 to VDR-RXR-DNA increases the average size of the complex but does not induce a large change in the overall SAXS profile: similar distinctive ‘humps’ around 1 nm^-1^ are observed in the SAXS curves for VDR-RXR-DNA alone and in complex with MED1 (Figure 2a,c). The probability distribution of real-space scattering pair distances, or *p*(*r*) profiles, reveals a similar shoulder around 60-80 Å for both complexes (Figure 2b). This indicates that the shape of the VDR-RXR-DNA in complex with MED1 remains similar to that of the DNA-bound heterodimer, where the DBDs are spatially separated from the LBDs. Comparison of the most representative *ab initio* models of VDR-RXR-DNA and VDR-RXR-DNA-MED1 complexes obtained with DAMMIN (Svergun, 1999) reveals overall shape similarities with two distinguishable domains (Figure 2c) indicating no major conformational change of the DNA-bound heterodimer upon MED1 interaction. For the coactivator complex, an additional electron density at the region occupied by both LBDs is visible indicating that the globular domain of MED1 is likely located on “top” of the heterodimer and is interacting with an extensive area within the VDR-RXR LBD heterodimer.

**Table 1.** SAXS parameters. Rg and Dmax as determined from Guinier plot or P(r) distribution.

| Sample | R<sub>g</sub>, Å<br>(from Guinier plot) | R<sub>g</sub>, Å<br>(from GNOM) | D<sub>max</sub>, Å<br>(from GNOM) |
| --- | --- | --- | --- |
| VDR-RXR-DNA-MED1 | 63.8 ± 2.3 | 63.6 ± 0.13 | 202 |
| VDR-RXR-DNA | 42.8 ± 0.14 | 43.3 ± 0.1 | 135 |
| MED1 | 41.5 ± 0.27 | 41.6 ± 0.12 | 140 |

**Figure 2.**
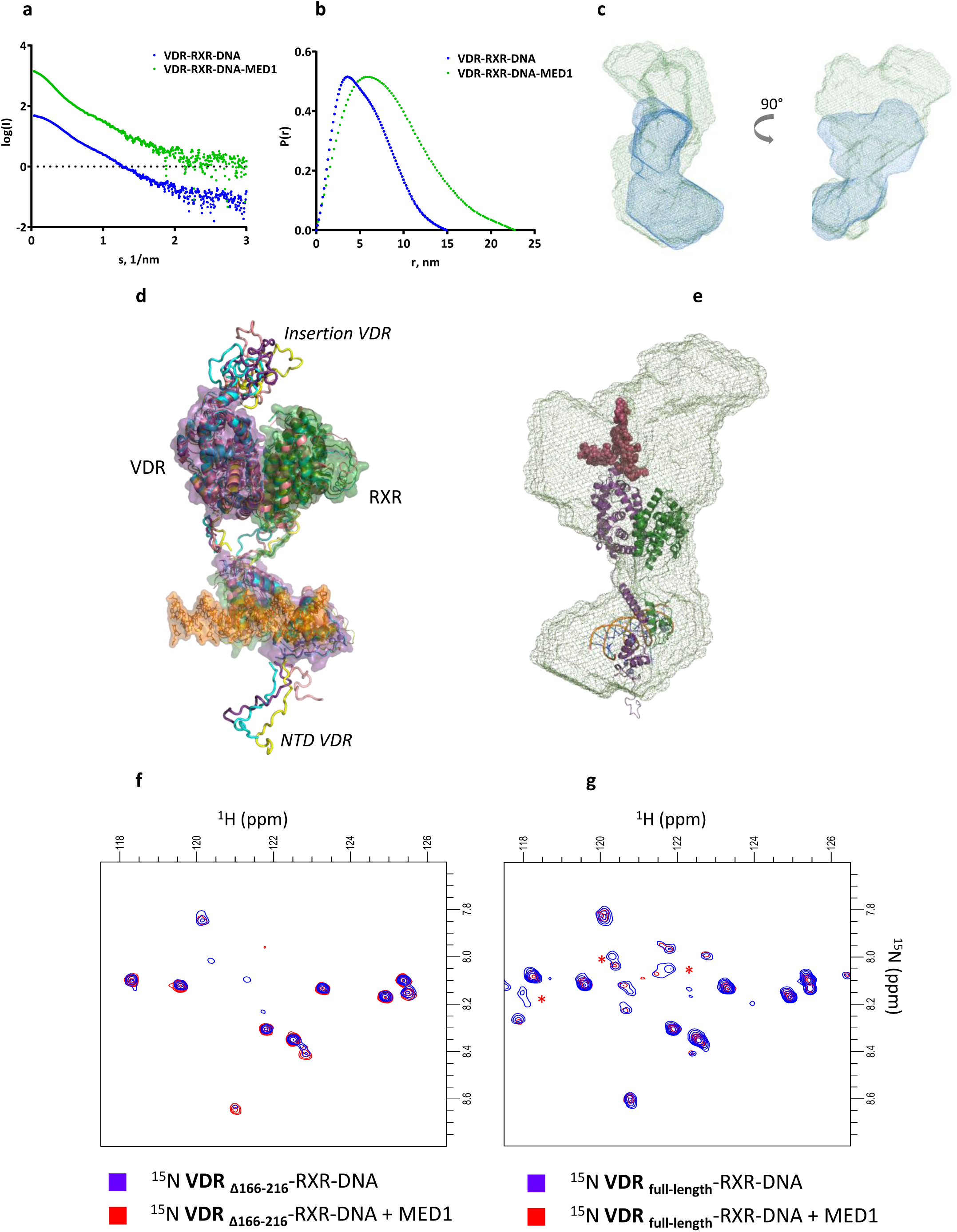
Solution structure of MED1 complex with VDR-RXR-rANF DR3. **(a)** Averaged SEC-SAXS data for VDR-RXR-DNA (blue) and VDR-RXR-DNA-MED1 (green) complexes. **(b)** Corresponding *p*(*r*) profiles calculated from the SAXS data. **(c)** Representative *ab initio* molecular envelopes for VDR-RXR-DNA-MED1 complex (green mesh) with superimposed ab initio molecular envelope of VDR-RXR-DNA (blue mesh). **(d)** SAXS-based ensemble models (4 conformations) of the VDR-RXR-DNA complex. **(e)** Best SAXS model of VDR-RXR-DNA fitted into the SAXS envelope of VDR-RXR-DNA-MED1. **(f)** ^1^H-^15^N HSQC spectra of the VDR-RXR-DNA complex lacking the VDR LBD insertion before (blue) and after addition of stoichiometric amount of MED1 (red), **(g)** ^1^H-^15^N HSQC spectra of the VDR-RXR-DNA complex harboring the full-length VDR before (blue) and after addition of stoichiometric amount of MED1 (red). The red stars indicate correlation peaks whose intensities are affected by the addition of MED1.

### VDR insertion domain modulates MED1 interaction with the VDR-RXR heterodimer

Whereas the *ab initio* SAXS envelope visualizes the overall shapes, the rigid-body refinement provides a model that reflects the overall distribution of conformers in solution and is not restricted to a particular low-energy conformation of the macromolecules. For the VDR-RXR-DNA complex, we built an ensemble of SAXS compatible models (Figure 2d) using the crystal structures of the DNA and ligand binding domains of VDR and RXR (Rochel et al., 2000; Egea et al., 2000; Shaffer and Gewirth, 2002) with missing regions (VDR NTD, hinges, VDR’s insertion) modeled as dummy residues. The insertion region in VDR LBD is a 50 amino acid domain specific for VDR and poorly conserved between VDR family members and disordered in context of the isolated LBD (Rochel et al., 2001). In the obtained refined models of VDR-RXR-DNA (Figure 2a) it occupies a defined region of space similar in all refined models suggesting that the insertion domain although possibly flexible is not totally disordered. Modelling also suggests that the VDR NTD (residues 1-23) is rather flexible, adopting various extended conformation in solution and not interacting with the DNA.

The best refined model docked into the SAXS envelope of VDR-RXR-DNA-MED1 (Figure 2e) reveals a proximity and possible overlap of the areas occupied by the VDR insertion domain and MED1, thus suggesting that this VDR region may be interacting with MED1. To further characterize the involvement of the disordered VDR insertion domain in MED1 association, a NMR analysis of N^15^-labelled VDR complexes was performed. As VDR exhibits two disordered regions, the short NTD and the insertion domain, we compared the ^1^H-^15^N HSQCs recorded for DNA bound heterodimer with VDR, either full-length or truncated of its insertion domain and with or without addition of MED1. The size of the complexes filters out signals from folded regions leaving only amide resonances from disordered regions belonging to the NTD and insertion in full-length LBD (Figure 2f,g). The addition of MED1 to VDR-RXR-DNA complex led to the specific disappearance of a small number of cross peaks specifically found in the full-length VDR indicating that the disordered insertion is involved, either directly or indirectly in the interaction with MED1.

### Roles of VDR and RXR AF2 in association with MED1

To analyze the roles of the VDR and RXR AF2 in interaction with MED1 (50-660) fragment, we mutated VDR and RXR and determined the impact of mutations on the association with MED1 by surface plasmon resonance. MED1 fails to interact to VDR-RXR when VDR H12 is deleted, or in presence of VDR antagonist ligand (ZK168281) (Figure 1c and Supplementary Figure 8), thus confirming that the main anchoring interaction are agonist-dependent and mediated through VDR AF-2, in agreement with previous studies showing that a synthetic peptide comprising classical NR-interacting LXXLL motif competes with MED1 for interaction with the VDR (Rachez et al., 2000). In contrast, deletion of RXR H12 does not prevent MED1 binding to VDR-RXR-DNA. A slightly increased MED1 binding is observed for VDR-RXRDeltaH12. Mutations of RXR residues forming a charge clamp for proper orientation and binding of the LXXLL motif of the coactivator (Gampe et al., 2000; Pogenberg et al. 2004), has no effect on MED1 interaction.

In addition, we investigated the impact of the binding of the peptide bearing the MED1 NR2 motif (residues 645-649) on the VDR-RXR-DNA complex using hydrogen-deuterium exchange coupled to mass spectrometry (HDX-MS). We observed that two regions of VDR corresponding to the NTD and the insertion domain exhibit fast H/D exchange rates (Supplementary Figure 6a) which is a typical phenomenon for highly flexible regions (Keppel et al., 2011), thus supporting SAXS results. Similar fast HDX rates were also observed for several regions of RXR (Supplementary Figure 6b), encompassing the hinge region, helix H1, H2, the C-terminus of H3, H11, the C-terminus of H12, and the region after H12. The comparison of relative fractional uptakes (RFU) of VDR-RXR-DNA and VDR-RXR-DNA-MED1 NR2 motif revealed that mainly VDR region 411-419 spanning the C-terminus of H11n and the N-terminus of the helix H12 was protected from H/D exchange upon NR2 motif binding (Figure 3 and Supplementary Figure 7a). These results are in agreement with the crystal structures of VDR LBD complexes with coactivator peptides (Vanhooke et al., 2004; Ciesielski et al., 2004), highlighting the role of VDR AF-2 in its interaction with MED1 NR2, through H3, H4 and H12 helices. On the contrary, no significant differences were detected for RXR upon MED1 NR2 motif binding (Figure 3 and Supplementary Figure 7b), supported by the low affinity of MED1 NR2 for RXR (Pogenberg et al., 2005; Stafslien et al., 2007). Taken together, our data show that the AF-2 of VDR but not RXR is directly involved in agonist-induced binding of MED1.

**Figure 3.**
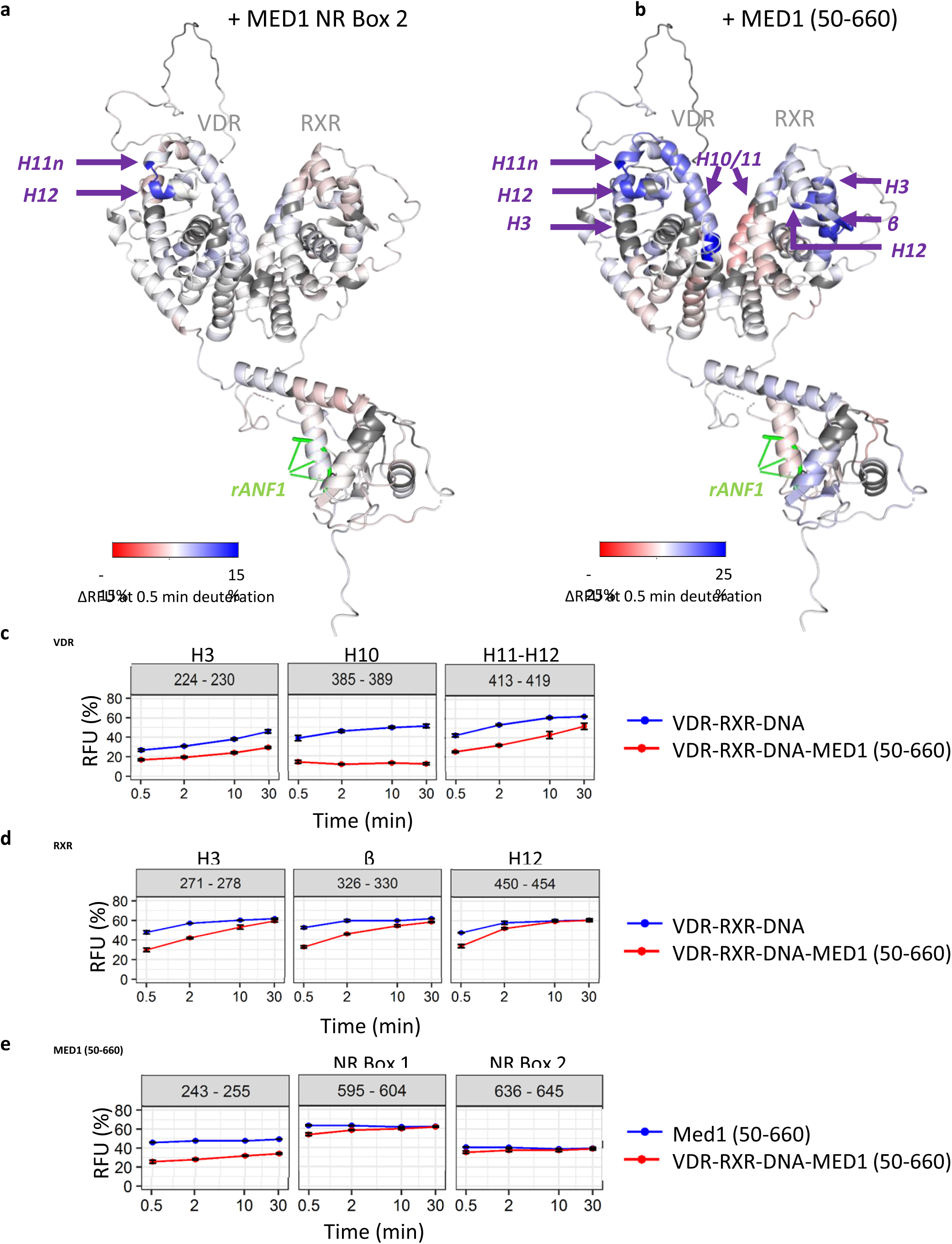
HDX-MS analysis of the VDR-RXR-DNA interaction with MED1. **(a)** Relative fractional uptake (RFU) differences between VDR-RXR-DNA and VDR-RXR-DNA-MED1 NR Box 2 complexes mapped on the representative SAXS model of the VDR-RXR-*rANF1* after 0.5 minutes of deuteration. **(b)** Relative fractional uptake (RFU) differences between VDR-RXR-DNA and VDR-RXR-DNA-MED1 (50-660) complexes mapped on the representative SAXS model of the VDR-RXR-*rANF1* after 0.5 minutes of deuteration. **(c)** Deuterium uptake of the peptides 224-230 (H3), 385-389 (H10) and 413-419 (H11-12) from VDR perturbed upon MED1 (50-660) binding plotted as a function of deuteration time. **(d)** Deuterium uptake of the peptides 271-278 (H3), 326-330 (β) and 450-454 (H12) from RXR perturbed upon MED1 (50-660) binding plotted as a function of deuteration time. **(e)** Deuterium uptake of selected MED1 peptides perturbed upon binding to VDR-RXR-DNA: 243-255 from the structured N-terminal part, 595-604 comprising Leu604 of the NR Box 1 and 636-645 comprising Leu645 from the NR Box 2.

### Effect of MED1 (50-660) binding on the VDR-RXR heterodimer

We next performed HDX-MS experiments with the larger construct of MED1 to determine the effect of binding on both the coactivator and the VDR-RXR heterodimer. HDX-MS of MED1 (50-660) fragment revealed that the protein is mostly structured (Supplementary Figure 9a), in agreement with the disorder prediction (Supplementary Figure 9b). Several MED1 regions exhibit fast H/D exchange rates and can be characterized as highly flexible, including residues 61-74, 161-174, 231-242, 276-288, 364-371 and 480-487 which could correspond to flexible loops between the secondary structure elements (Supplementary Figure 9c). MED1 RID is also highly flexible, in agreement with our previous data (Rochel et al., 2011).

We compared HDX-MS RFU values of VDR and RXR in VDR-RXR-DNA complex with those in VDR-RXR-DNA-MED1 (50-660) state to characterize their conformational dynamics upon MED1 binding. As expected, C-terminus of H11n and N-terminus of H12 of VDR (protected upon MED1 NR2 binding) were similarly affected upon MED1 (50-660) binding (Figure 3 and Supplementary Figure 10a). Additional regions of VDR were protected, including helices H3, the C-terminus of H5, H6, H7, H10 and H11. These regions are spatially close to the VDR-AF2 domain (Figure 3b), revealing a higher protection effect on this region upon MED1 (50-660) binding compared to the binding of MED1 NR2 motif.

Interestingly, several protected regions of RXR LBD were identified upon MED1 (50-660) binding, corresponding to helices H3, H5 and the β-strand, H11 and the N-terminus of H12 (Figure 3 and Supplementary 10b). These regions are spatially close to each other and are located on “top” of the heterodimer LBDs supporting the SAXS models indicating that this region creates a large MED1 interaction surface. Of note, region 419-429 of RXR, covering helix H10 and comprising the heterodimerization interface, shows deprotection upon MED1 binding at shorter time points suggesting a conformational change leading to its higher flexibility (Figure 3b and Supplementary Figure 9b).

MED1 (50-660) is also protected upon formation of the complex with VDR-RXR-DNA (Supplementary Figure 11). Residues 243-255 of the structured N-terminal domain of MED1 are particularly protected upon VDR-RXR-DNA binding (Figure 3e). Other affected regions are 74-107, 123-150, 191-201, 509-527 as well as regions 560-604, 636-645 and 649-657 situated within the unstructured part of MED1 comprising RID. Regions 560-604 and 636-645 encompass the first leucine residues of both NR boxes 1 and 2 of MED1 RID domain respectively (Leu604 and Leu645), suggesting a stabilization of both motifs upon NR binding.

All together, these results show that MED1 (50-660) binding affects extended regions within the liganded VDR-RXR LBD heterodimer, in contrast to the MED1 NR2 motif stabilizing solely the C-terminus of VDR. In addition to the VDR coactivator cleft and H12 strongly stabilized upon MED1 binding, several RXR regions including AF-2 and H5/β-strand are affected indicating role of RXR in establishing a specific association with the coactivator.

### Investigation of the interaction surface between MED1 and VDR-RXR heterodimer

To determine molecular constraints and amino acids of VDR-RXR and MED1 located in close proximity, we next performed crosslinking mass spectrometry experiments (XL-MS) on VDR-RXR-DNA-MED1 (50-660) complex using DSBU (spacer arm 12.5 Å) (Müller et al. 2010) and a C2-arm version of the DSBU (spacer arm 6.2 Å) (see material and methods). Both crosslinkers are MS-cleavable, allowing more confident MS/MS identification and validation thanks to the detection of characteristic doublet peaks along with the peptide backbone fragments (Arlt et al., 2016). Both DSBU crosslinker versions target primary amines as well as hydroxyl groups and can bridge residues with Cα-Cα distances up to 26-30 Å and 20-24 Å for DSBU and the C2-arm, respectively (Merkley et al., 2014). We identified 42 intra- and inter-crosslinked peptides: 11 crosslinks intra-RXR, 12 intra-VDR, 6 intra-MED1, 12 inter-VDR-RXR and 1 inter-MED1-RXR (Figure 4 and Supplementary S12a). Twelve crosslinks involving VDR and RXR were found in the proximity to or within the DBDs when mapped on the VDR-RXR heterodimer model (Orlov et al., 2012) and with a Cα-Cα distance below the cut-off mentioned, increasing confidence in our XL-MS approach (Supplementary Figure 12b). Interestingly, an inter-protein crosslink involving Thr236 of MED1 and Lys321 of RXR was identified. Lys321 is located in the β-strand between H5 and H6 of RXR LBD (Figure 4b) on “top” of the LBD heterodimer and within the region protected from the H/D exchange upon MED1 binding correlating with the SAXS and HDX-MS results. Taken together, our data suggests that although RXR H12 is not required for MED1 recruitment, RXR LBD possesses an extensive surface directly interacting with MED1.

**Figure 4.**
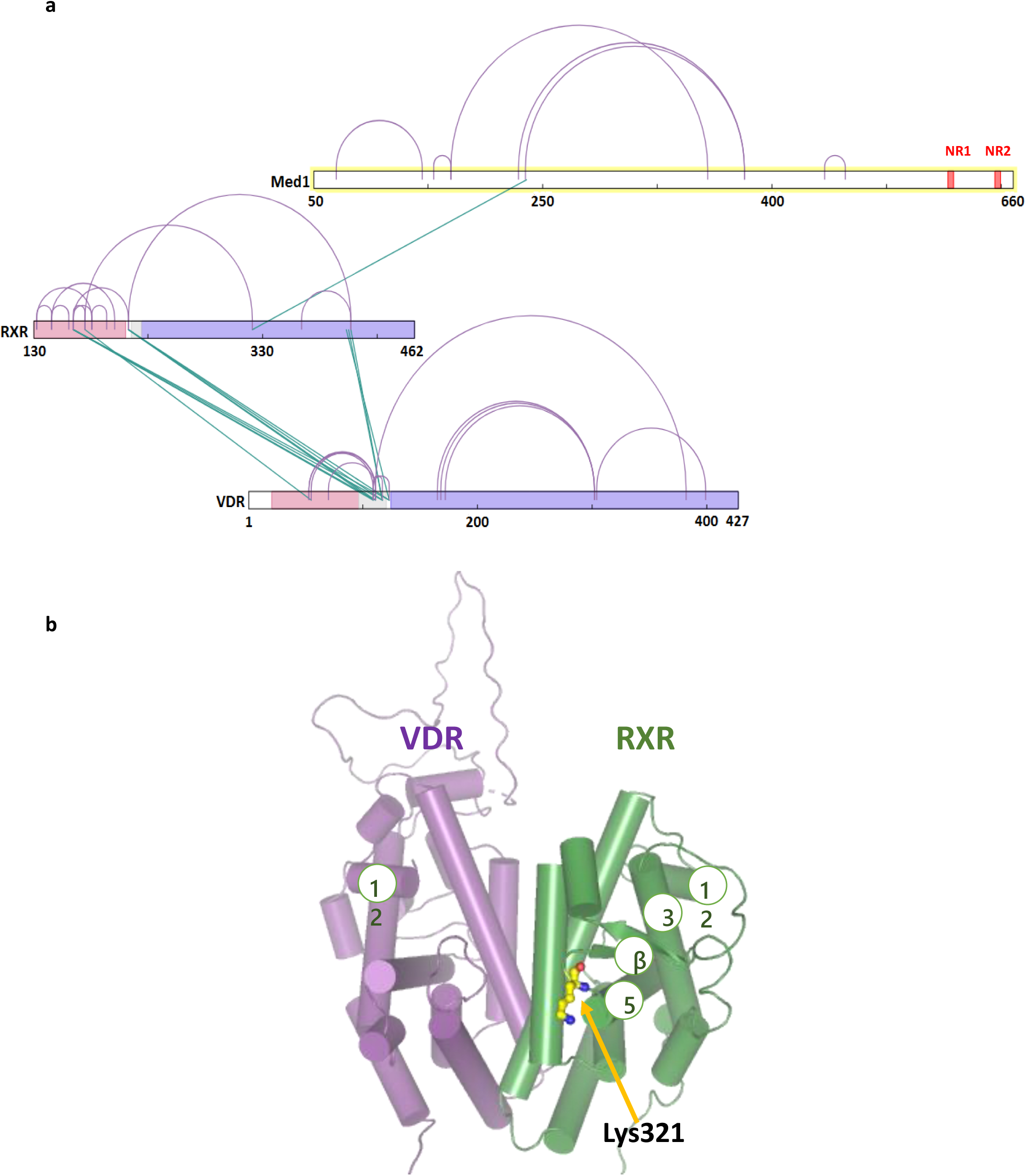
Crosslinked sites observed for the VDR-RXR-DNA-MED1(50-660) complex. **(a)** Identified VDR-RXR-MED1 (50-660) crosslinks representation. Red: NR boxes 1 and 2 of MED1; purple: LBD region; grey: hinge region; pink: DBD region for each nuclear receptor, VDR and RXR. Blue lines represent inter-protein crosslinks and purple curved lines intra-protein crosslinks. **(b)** Representation of Lys321 of RXR, identifies as crosslinked to Thr236 of MED1, within the VDR-RXR LBD heterodimer.

## Discussion

While structural requirements for ligand-dependent binding of classical coactivator motifs to NRs have been fully elucidated (reviewed in Weikum et al., 2018), the molecular mechanism of co-activation remains poorly understood. In particular, studies providing structural insights into recognition of full-length or truncated coactivators by full NR complexes remain extremely scarce (Yi et al., 2015; Yi et al., 2017; Rochel et al., 2011; Zhang et al., 2011). Here, we describe novel structural data on the complex formed between the agonist- and DNA-bound VDR-RXR heterodimer and a large fragment of the classical nuclear receptor coactivator MED1.

MED1 interacts with NRs through a disordered RID domain that contains two LXXLL motifs (Rachez et al., 2000). Early mutagenesis studies revealed that in case of the VDR-RXR heterodimer VDR binds with high affinity the second LXXLL motif of MED1 but not the NR1 box, and RXR shows only weak binding to both motifs (Ren et al. 2000; Yang et al., 2000; Malik et al. 2004). In addition, mutations in the VDR’s charge clamp render VDR inactive, confirming that interaction with coactivators is crucial for its activity (Jimenez-Lara and Aranda 1999). Our results confirm that MED1 NR2-VDR interaction is driving the complex formation. However, by using a larger fragment of MED1, MED1 (50-660), we have also demonstrated that other VDR-RXR regions outside the VDR AF2 as well as MED1 regions other than RID modulate the association and form an extended interaction surface.

Both LXXLL motifs of MED1, as well as proper spacing between the motifs, were reported to be required for optimal MED1 binding to DNA-bound VDR-RXR heterodimer (Ren et al. 2000). It was suggested that each LXXLL motif is recognized by VDR and RXR coactivator binding clefts simultaneously when the complex with MED1 is formed. In this case the 35 amino acid spacer between the two LXXLL motifs has to clasp around the LBDs as the coactivator binding sites are located on the opposite sites of the heterodimer. However, we have previously shown that the MED1 RID binds to the VDR-RXR heterodimer asymmetrically and remains flexible (Rochel et al., 2011). Interestingly, in the present study, we observed the perturbation of both LXXLL motifs of MED1 (50-660) upon formation of the complex with VDR-RXR, suggesting that while the NR1 box is not accommodated within the classical coactivator binding site, it could be either interacting with an alternative site of the receptors or stabilized allosterically. Similar observations have previously been made for the three NR boxes of the SRC-2 RID binding to PPARG-RXR (de Vera et al., 2017). Alternative interaction with AR AF-1 was previously described for two noncanonical α-helical motifs of MED1 located between residues 505 and 537 and in proximity to the NR box 1 (Jin et al., 2011).

Here we demonstrate that, in addition to RID, the structured N-terminal domain of MED1 is alsoaffected upon binding to VDR-RXR and is likely interacting with both VDR and RXR LBDs. In particular, MED1 region 243-255 is largely stabilized in the complex with the receptor heterodimer. Neighboring Thr236 of MED1 was identified within inter-MED1-RXR crosslink, suggesting that this MED1 region is in physical proximity to the RXR β-strand which, in turn, is also perturbed upon the interaction. As N-terminal region of MED1 is involved in important downstream interactions such as incorporation into Mediator (Ge et al., 2008) and recruitment of alternative cofactors to enhancer-bound NRs (Iida et al., 2015), it is tempting to speculate that by creating an extended surface for MED1 accommodation and altering its conformation, RXR could be important for achieving optimal MED1-mediated transcription activation.

Among other novel MED1-interacting regions within the VDR-RXR heterodimer is the flexible insertion domain in the VDR LBD located between H1 and H3. By using ^15^N NMR, we show that it undergoes conformational changes upon interaction with MED1. This effect is not seen in the HDX-MS experiment; however the observed difference could be attributed to a different temporal resolution of the two methods. While the VDR insertion domain does not play a major role in receptor selectivity for 1,25D3 (Rochel et al., 2001) nor bile acids (Krasowski et al., 2011), previously published data suggests that it has a functional role in regulation of VDR signaling. Several phosphorylations modulating VDR transcriptional activity, S182 and S208, have been identified within this region (Zenata and Vrzal, 2017). Interestingly, S208 phosphorylation has previously been reported to enhance VDR interaction with MED1 but not with the SRC1 coactivator (Bareletta et al., 2002: Arriagada et al., 2007).

In this study we used the full VDR-RXR heterodimer in agonist-bound form, where both VDR and RXR ligands were present. Hypothetically, each receptor can recruit a coactivator, however, we demonstrate that only one molecule of MED1 is recruited by the VDR-RXR heterodimer, suggesting that H12 of RXR is not essential for the complex formation. Indeed, truncation of RXR H12 and mutations in RXR charge clamp do not prevent MED1 interaction, and binding of a peptide comprising MED1 NR2 does not induce any change in RXR as seen by HDX-MS. However, using HDX-MS, we identified H3, H11 and the N-terminus of H12 comprising the AF-2 among RXR regions largely stabilized upon MED1 binding. The observed stabilization event could originate from direct non-canonical interaction with MED1 as well as from a distal allosteric effect.

As MED1 is a classical NR coactivator, it is recognized by NRs similarly to other coregulators, e.g. SRCs, and their binding sites overlap. Interestingly, significant differences could be observed between the VDR-RXR complex with MED1 described here and the analogue complexes with the SRCs. In presence of 1,25D3 only RXR AF2 within the full VDR-RXR complex was insensitive to the binding of SRC1 RID, indicating that it is primarily associated to VDR AF2 (Zheng et al. 2015). However, RXR and its ligand modulate the interaction and in the presence of both VDR and RXR agonists SRC1 RID binding has been shown to stabilize VDR AF2 as well as RXR H3 (227-273) and H10-H11 (433-451) (Zhang et al. 2011). Differences in binding mode between SRCs and MED1 were previously suggested for ER where a single mutation differently affects MED1 and SRC2 RID interaction (Acevedo et al., 2004), or for PPAR-RXR where RXR ligand only induces SRCs but not MED1 binding to RXR (Yang et al. 2000). Such differences in the binding modes could serve as molecular determinants of how the NRs discriminate between coactivators and sequentially recruit them.

This work contributes to a growing number of studies revealing the complexity of coactivator binding to NRs. Extended characterization of allosteric mechanism within large NR coregulator complexes should increase the potential of novel targets for drug design and discovery programs.

## Acknowledgements

We thank the staff of BM29 at ESRF for assistance in using the beamline. We thank the IGBMC cell culture service, Isabelle Kolb-Cheynel and Nathalie Troffer-Charlier (IGBMC) for insect cell production, Pascal Eberling (IGBMC) for peptide synthesis, Pierre Poussin-Courmontagne (IGBMC) for assistance with SEC-MALLS and Catherine Birck (IGBMC) for help with AUC data collection.

## Supporting Information

Additional supplementary methods and figures can be found in SI appendix.

