## Supplementary Data for "Molecular determinants of MED1 interaction with the DNA bound VDR-RXR heterodimer"

#Co-first authors

🖂 Correspondence to Anna Y. Belorusova or Natacha Rochel

### Supplementary Methods

**Isothermal titration calorimetry.** Measurements were performed at 15°C on a MicroCal ITC200 (MicroCal). Oligonucleotides and purified VDR-RXR were dialyzed extensively against the buffer containing 20 mM HEPES pH 7.5, 100 mM NaCl, 5% Glycerol, 1 mM TCEP overnight. Direct titration experiments were performed as follows: 2 µL aliquots of DR3 at 75–150 µM were injected into a 7.5 – 15 µM VDR-RXR solution in a 200 µL sample cell. The duration of each injection was 4 s with a delay between injections of 120 s. ITC titration curves were analyzed using the software Origin 7.0 (OriginLab). Standard free energies of binding and entropic contributions were obtained, respectively, as $\Delta G = -RT\ln(Ka)$ and $T\Delta S = \Delta H - \Delta G$, where the association constant Ka and enthalpy change ΔH values were derived from ITC curve fitting.

**Reporter gene assay**

HEK293 EBNA cells were plated into 24-well plates at 105 cells per well and grown overnight in Dulbecco's modified Eagle's medium (DMEM) supplemented with 10% charcoal- treated fetal bovine serum (FCS) and 40 µg/mL gentamycin. At 80% confluence cells were transfected with 1 µg of pDNA per well using jetPEI (Polyplus transfection). Transfection was performed according to the manufacturer's instructions. The transfection mix consisted of the cells were transfected with 150ng of the expression plasmid pSG5-hVDR (1-427), 150 ng of the pSG5-hRXRα (1-462), 150 ng of the reporter plasmid containing the DR3-type *rANF1* vitamin D response element fused to the tk promoter, 3ng of the pRL plasmid (Promega) containing the Renilla luciferase gene (transfection and cellviability control), and 497 ng of the carrier plasmid pBlueScript (Stratagene). Eight hours post transfection, 1,25D3 or vehicle (ethanol) were added. Cells were harvested after eighteen hours of incubation with the ligand. The amounts of reporter gene product (firefly luciferase) and constitutively expressed Renilla luciferase produced in the cells were measured using Dual-Luciferase® Reporter Assay System (Promega) on a luminometer plate reader LB96P (Berthold Technologies). Luminescence of firefly luciferase values was normalized to the Renilla luciferase activity. Luciferase activities were expressed as relative light units (RLU) intensity. Assays were performed in triplicate for at least two independent experiments. For every triplicate, the mean and the standard error of the mean were calculated.

**Supplementary Figures Legend**

**Supplementary Figure 1. (a)** Structural organization of hMED1. **(b)** Disorder prediction for MED1 obtained by RONN (http://www.strubi.ox.ac.uk/RONN).

**Supplementary Figure 2. VDR-RXR-DNA-MED1 complex. (a)** ITC binding isotherm (upper panel) and fit to the binding curve (lower panel) for *rANF1* DR3 binding to the VDR-RXR. **(b)** Transactivation assay in HEK293 EBNA cells co-transfected with pDNAs encoding full-length VDR and luciferase cloned with two copies of the *rANF1* VDRE. Luciferase activity was measured after 24 hours of the cells treatment with increasing amounts of 1,25D3. **(c)** Gel retardation in TBE. 6% acrylamide gel stained with Coomassie Blue. **(d)** Overlay of gel filtration chromatograms for VDR-RXR-DNA, MED1 and their mix, and analysis of gel filtration fractions by a 10% SDS-PAGE gel. Fractions of interest are framed.

**Supplementary Figure 3. Sedimentation profiles of the MED1 binding reaction.** The MED1:VDR-RXR-DNA ratio is indicated above each box. Upper panels: raw sedimentation data (dots) and fitted c(s) distributions. Lower panels: residuals of the c(s) fit. Early data/fits/residuals are colored violet, and later data are colored according to the rainbow.

**Supplementary Figure 4. On-line SAXS coupled with SEC.** Gel filtration was performed on the Superdex S200 Increase column (GE Healthcare). (**a**) Elution profiles. (**b**) Plots of total scattering (upper panel) and radius of gyration (lower panel) vs. frame.

**Supplementary Figure 5. SAXS analysis of MED1 (50-660). (a)** SAXS profile after an on-line GF separation together with the corresponding fit of the theoretical data for the refined model. **(b)** *p*(*r*) profile calculated from the SAXS data. **(c)** Kratky plot.

**Supplementary Figure 6.** Relative fractional uptakes of VDR **(a)** and RXR **(b)** represented for VDR-RXR-DNA and VDR-RXR-DNA-MED1 NR Box 2 states at all deuteration times (0.5, 2, 10 and 30 minutes). Red framed peptides correspond to highly flexible regions of these proteins, presenting fast exchange rates. Some of RFU plots of these regions are represented.

**Supplementary Figure 7.** Relative fractional uptake difference plots represented for VDR **(a)** and RXR **(b)** profiles displaying change in HDX upon binding of MED1 NR Box 2 peptide. RFU differences are depicted for 0.5, 2, 10 and 30 minutes of deuteration. Framed peptides represent the most impacted regions of VDR upon NR2 motif binding presenting a statistical significance ($p$<0.01, Wald Test, MEMHDX software) for the magnitude of the difference. Among them, blue highlighted peptides present RFU differences above 5% while grey highlighted peptides present RFU differences below 5%.

**Supplementary Figure 8**. **Gel filtration profiles. (a)** Overlay of gel filtrations chromatograms of VDR-RXR-DNA-MED1 and VDRΔH12-RXR-DNA-MED1 mix. **(b)** Overlay of gel filtrations chromatograms of VDR-RXR-DNA-MED1 mix in presence of 1,25D3 and 9cisRA, or ZK168281 alone or ZK168281 and 9cis RA.

**Supplementary Figure 9. (a)** Relative fractional uptake differences plots of MED1 (50-660) measured after 0.5, 2, 10 and 30 minutes of deuteration. (**b**) Disorder prediction for MED1. (**c**) Secondary structure prediction of MED1 (50-660) where α-helices are indicated in red, β-strands in purple and random coil in green.

**Supplementary Figure 10**. Relative fractional uptake difference plots of VDR **(a)** and RXR **(b)** in VDR-RXR-DNA and VDR-RXR-DNA-MED1 (50-660) states measured after 0.5, 2, 10 and 30 minutes of deuteration. Framed peptides represent the most impacted regions of VDR and RXR upon MED1 (50-660) binding presenting a statistical significance ($p$<0.01, Wald Test, MEMHDX software) for the magnitude of the difference. Among them, blue highlighted peptides present RFU differences above 5% while grey highlighted peptides present RFU differences below 5%.

**Supplementary Figure 11**. HDX-MS characterization of MED1 (50-660). **(a)** Heat map representation of MED1 (50-660) where RFU differences between MED1 (50-660) and VDR-RXR-DNA-MED1 (50-660) complex are depicted for 0.5, 2, 10 and 30 minutes deuteration times with a color scheme representing RFU differences (-/+ 15% difference range). **(b)** Relative fractional uptake difference plot of MED1 (50-660) in free and bound to VDR-RXR-DNA states after 0.5, 2, 10 and 30 minutes of deuterations. Framed peptides represent the most impacted regions of MED1 (50-660) upon VDR-RXR-DNA binding presenting a statistical significance ($p<0.01$, Wald Test, MEMHDX software) for the magnitude of the difference. Among them, blue highlighted peptides present RFU differences above 5% of RFU while grey highlighted peptides present RFU differences below 5% of RFU.

**Supplementary Figure 12**. VDR-RXR-DNA-MED1 crosslink experiment. **(a)** Table summarizing all identified crosslinked sites (inter and intra) for the two used crosslinking agents. Cα-Cα distances are indicated for all identified VDR-RXR inter crosslinked peptides and where no distance was observed over the cut-off distance of each crosslinker (26-30Å and 20-24Å for DSBU and C2-arm version respectively). **(b)** Cα-Cα distances for inter VDR-RXR identified crosslinks are represented on the heterodimer PyMOL structure.
