## Supplementary Figures for "Molecular determinants of MED1 interaction with the DNA bound VDR-RXR heterodimer"

Figure S1

a

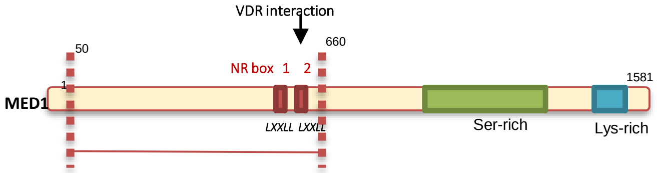

VDR interaction
50
660
NR box 1 2
1581
1
MED1
LXXLL LXXLL
Ser-rich
Lys-rich


b

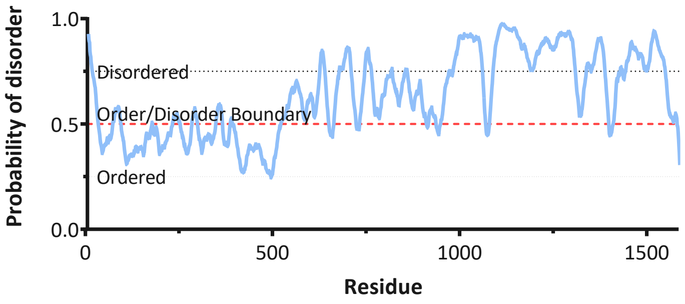

Probability of disorder
1.0
0.5
0.0
Disordered
Order/Disorder Boundary
Ordered
0
500
1000
1500
Residue

**Figure S2**

a

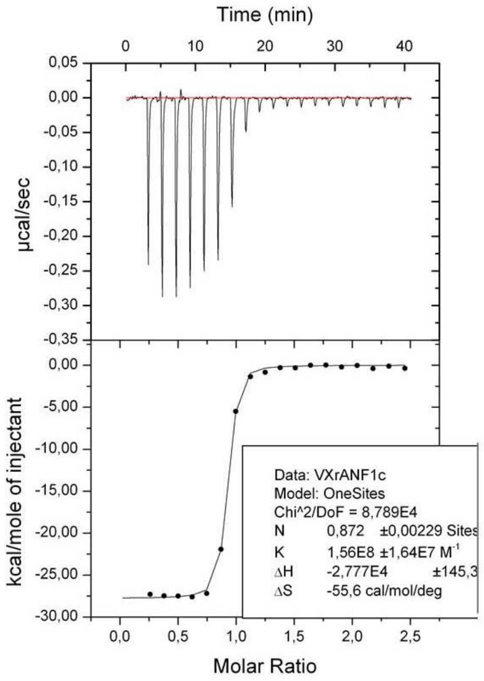


b

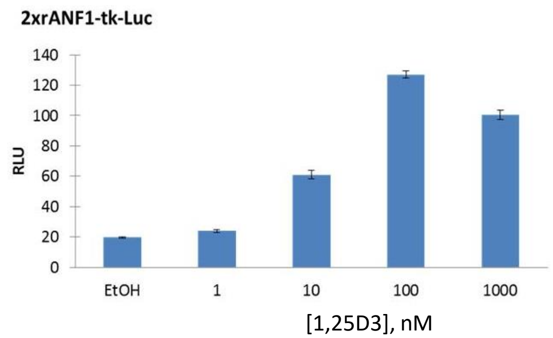


c

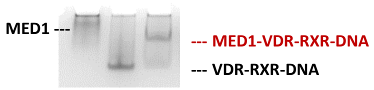


d

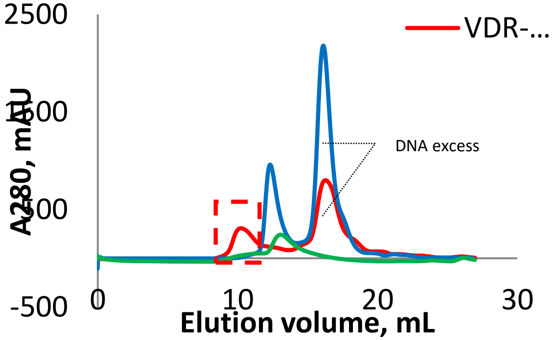


e

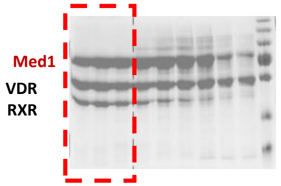

**Figure S3**

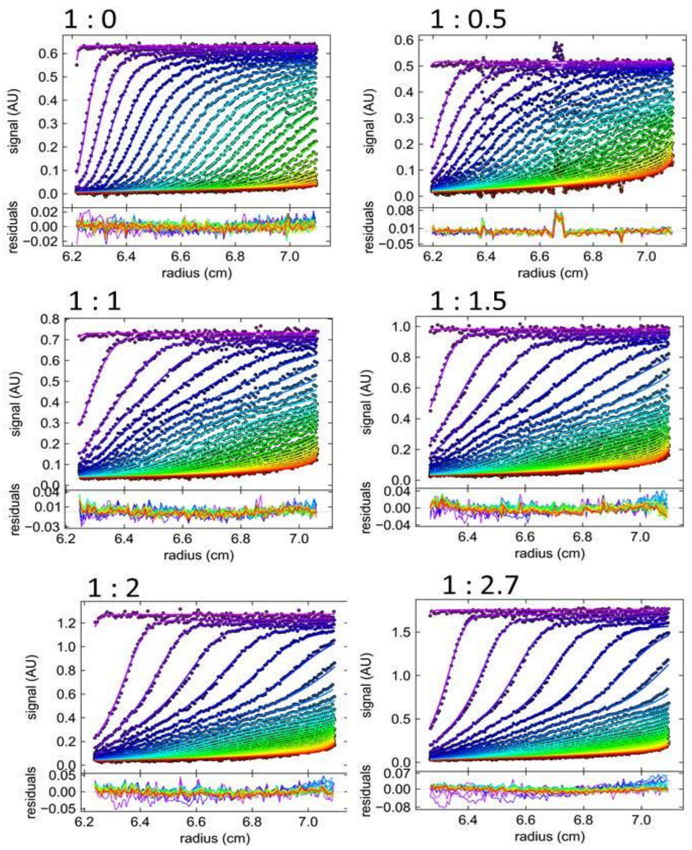

### Figure S4

a

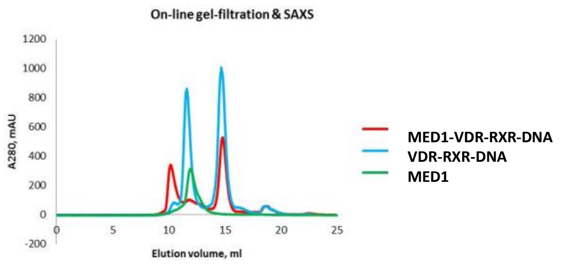
On-line gel-filtration & SAXS
A280, mAU
Elution volume, ml
MED1-VDR-RXR-DNA
VDR-RXR-DNA
MED1

b

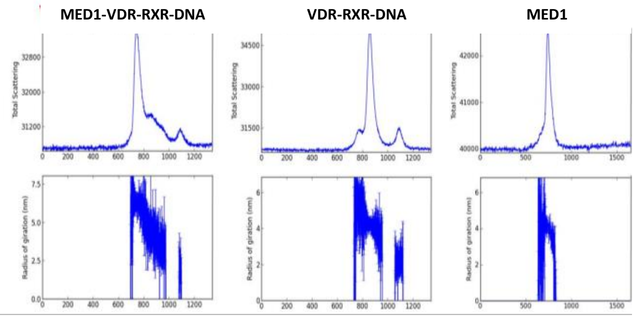
MED1-VDR-RXR-DNA
VDR-RXR-DNA
MED1
Total Scattering
Radius of gyration (nm)

**Figure S5**

a

b

c

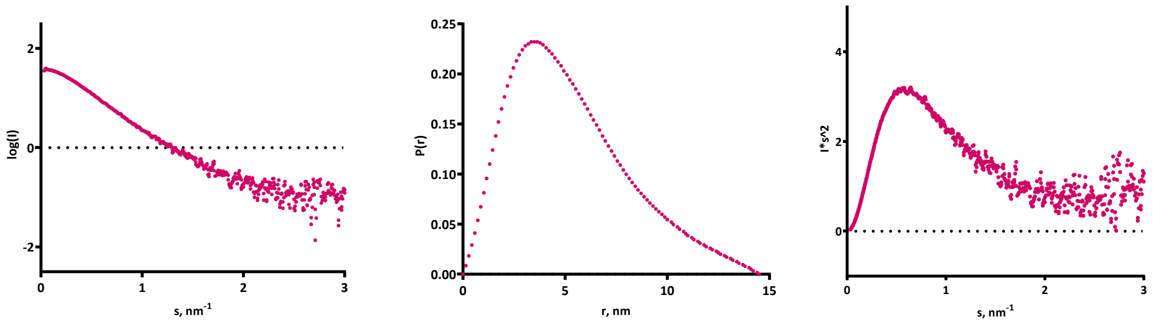

### Figure S6

**a** **RFU of VDR-RXR-*rANF1* *vs* VDR-RXR-*rANF1*-NR2 : VDR profile**

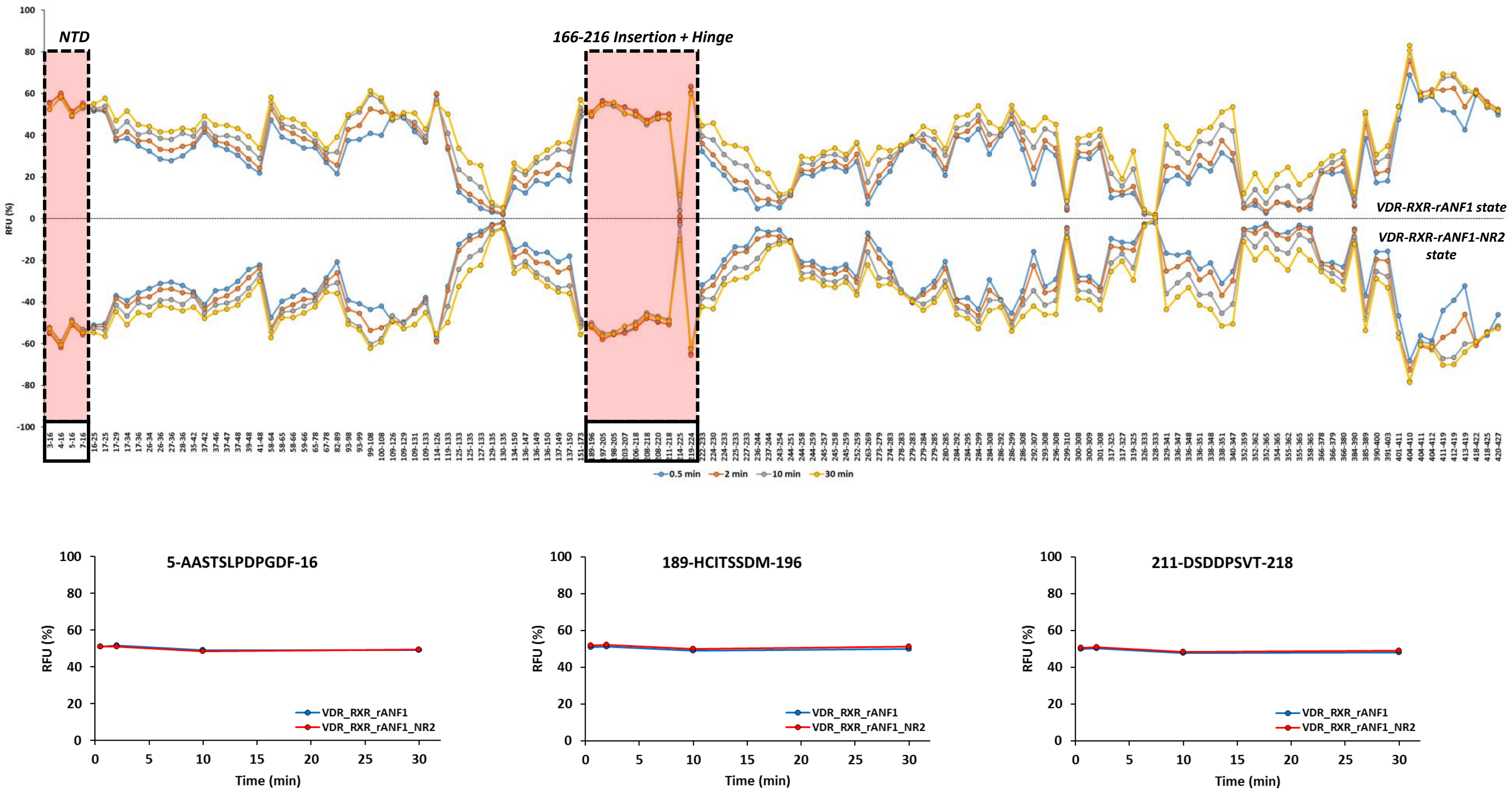

**Figure S6**

**b** **RFU of VDR-RXR-*rANF1* *vs* VDR-RXR-*rANF1*-NR2 : RXR profile**

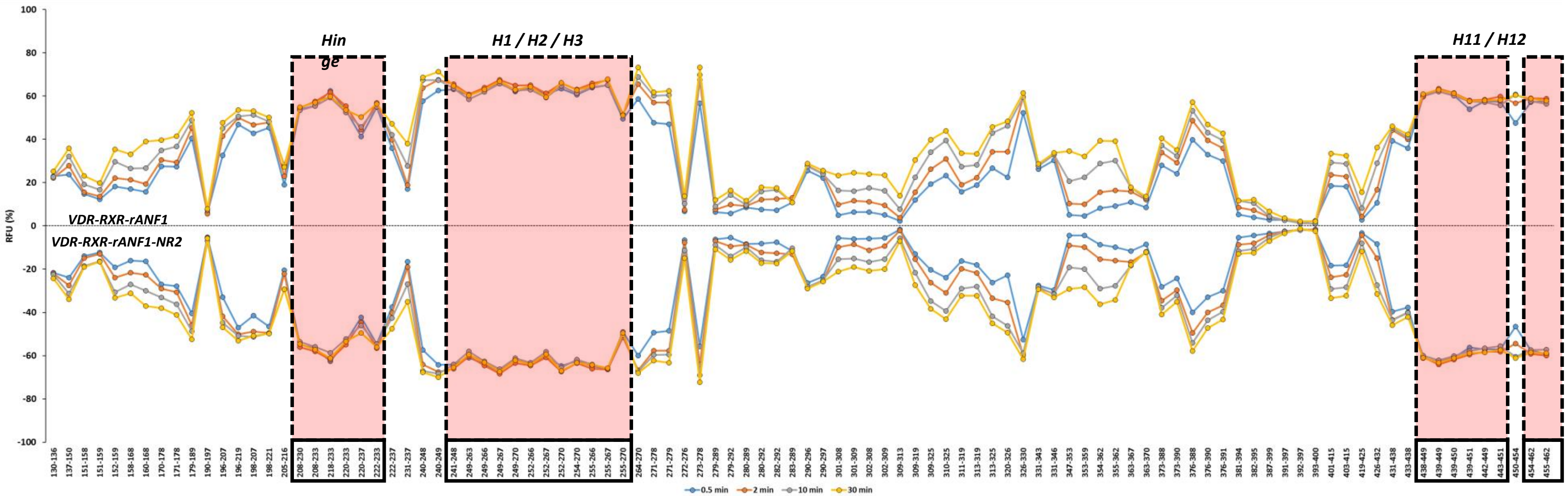


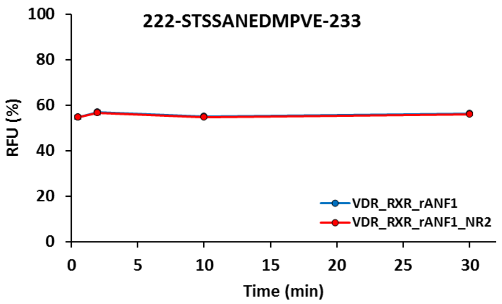


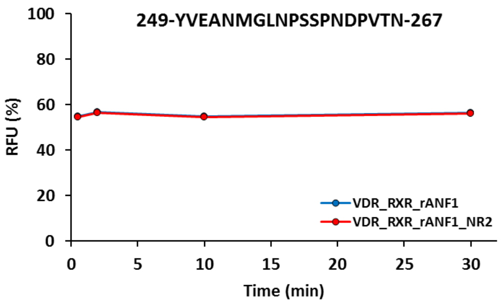


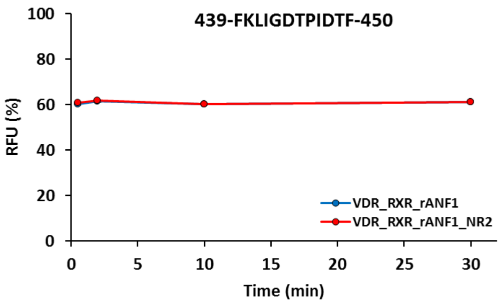

**Figure S7**

**a** **RFU Difference of VDR-RXR-*rANF1* *vs* VDR-RXR-*rANF1*-NR2 : VDR profile**

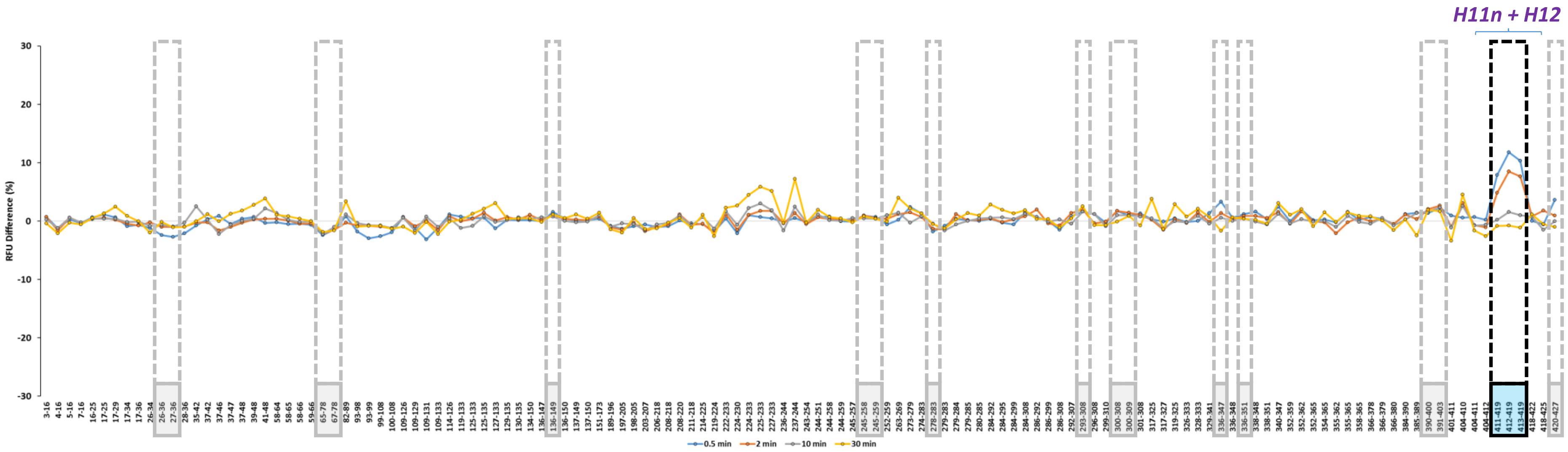


**b** **RFU Difference of VDR-RXR-*rANF1* *vs* VDR-RXR-*rANF1*-NR2 : RXR profile**

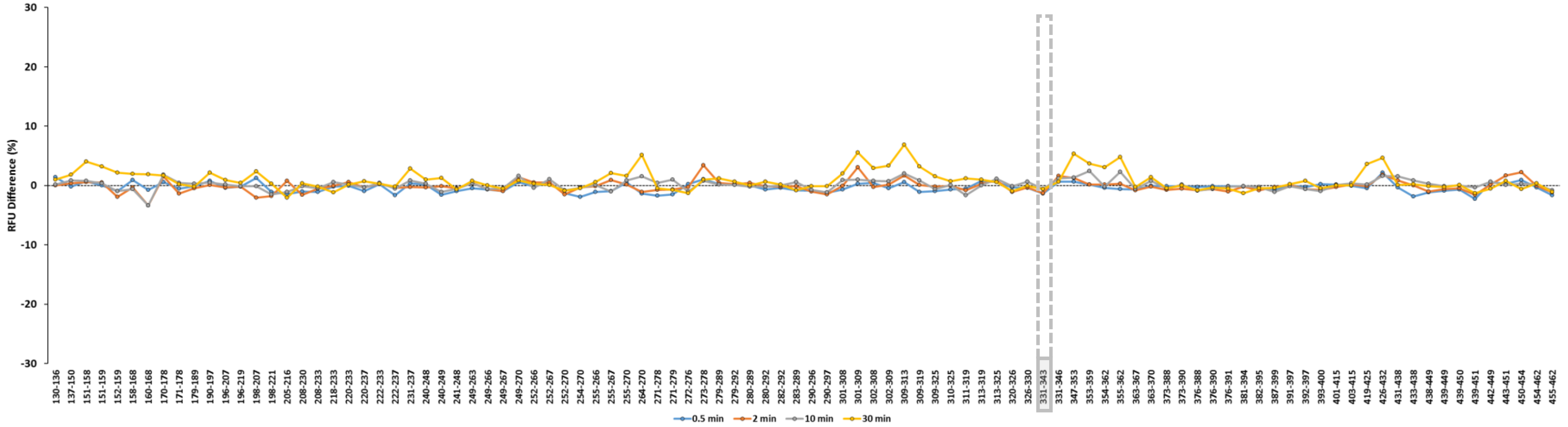

**Figure S8**

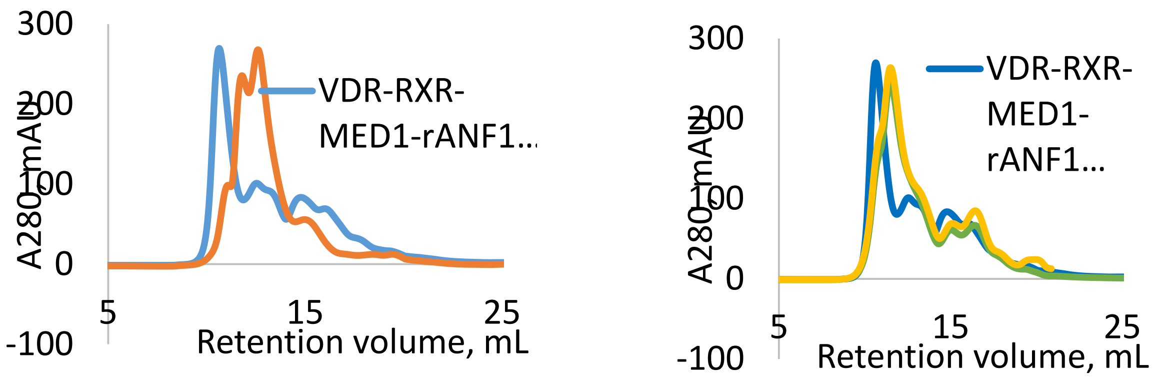
300
200
100
0
-100
A280, mAU
5
15
25
Retention volume, mL
VDR-RXR-
MED1-rANF1...
300
200
100
0
-100
A280, mAU
5
15
25
Retention volume, mL
VDR-RXR-
MED1-
rANF1...

**Figure S9**

**a**

**RFU of MED1**

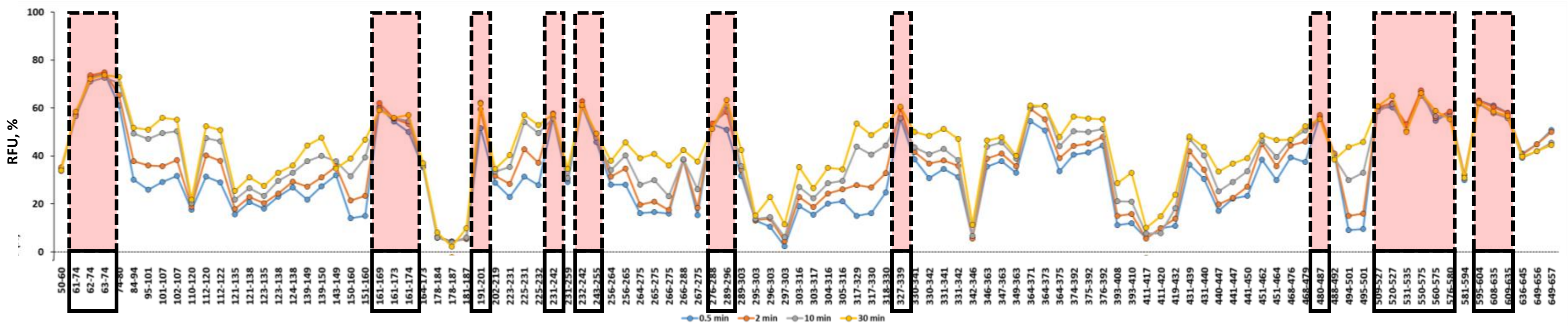


**b**

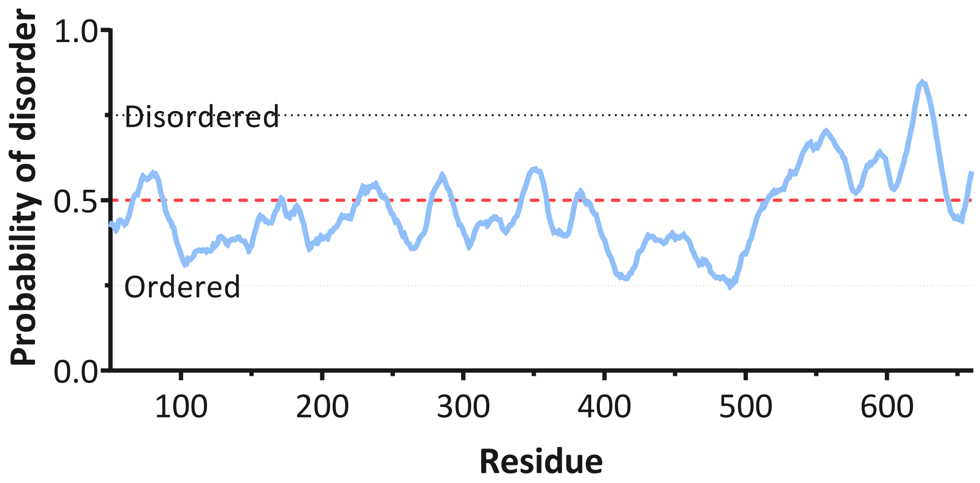


**c**

MSSGGHQHLV SCLETLQKAL KVTSLPAMTD RLESIARQNG
LGSHLSASGT ECYITSDMFY VEVQLDPAGQ LCDVKVAHHG
ENPVSCPELV QQLREKNFDE FSKHLKGLVN LYNLPGDNKL
KTKMYLALQS LEQDLSKMAI MYWKATNAGP LDKILHGSVG
YLTPRSGGHL MNLKYYVSPS DLLDDKTASP IILHENNVSR
SLGMNASVTI EGTSAVYKLP IAPLIMGSHP VDNKWTPSFS
SITSANSVDL PACFFLKFPQ PIPVSRAFVQ KLQNCTGIPL
FETQPTYAPL YELITQFELS KDPDPIPLNH NMRFYAALPG
QQHCYFLNKD APLPDGRSLQ GTLVSKITFQ HPGRVPLILN
LIRHQVAYNT LIGSCVKRTI LKEDSPGLLQ FEVCPLSESR
FSVSFQHPVN DSLVCVVMDV QDSTHVSCKL YKGLSDALIC
TDDFIAKVVQ RCMSIPVTMR AIRRKAETIQ ADTPALSLIA
ETVEDMVKKN LPPASSPGYG MTTGNNPMSG TTTPTNTFPG
GPITTLFNMS MSIKDRHESV GHGEDFSKVS QNPILTSLLQ
ITGNGGSTIG SSPTPPHHTP PPVSSMAGNT KNHPMLMNLL
KDNPAQDFST L

**Figure S10**

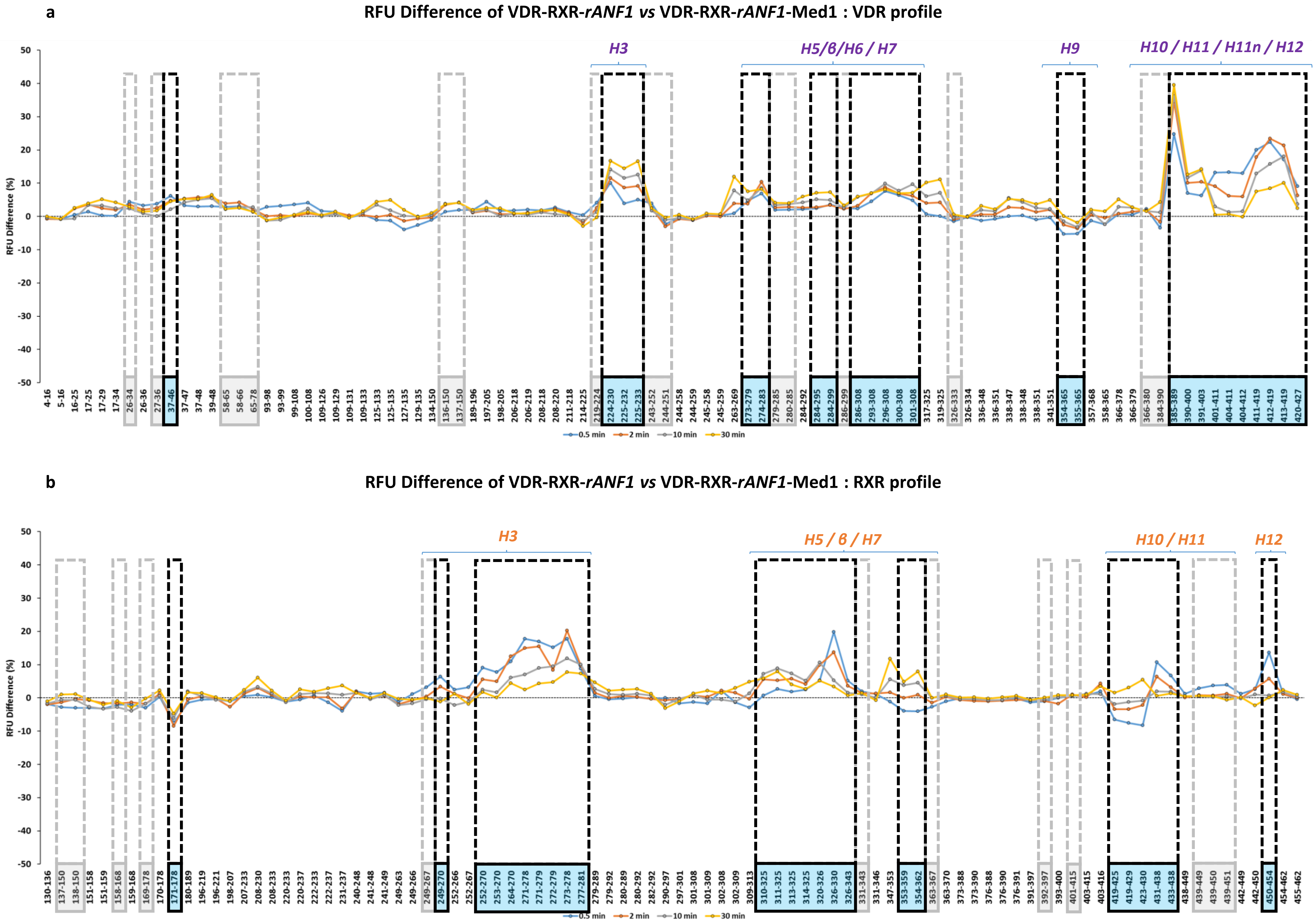
a
RFU Difference of VDR-RXR-rANF1 vs VDR-RXR-rANF1-Med1 : VDR profile
RFU Difference (%)
H3
H5/β/H6 / H7
H9
H10 / H11 / H11n / H12
0.5 min
2 min
10 min
30 min
b
RFU Difference of VDR-RXR-rANF1 vs VDR-RXR-rANF1-Med1 : RXR profile
RFU Difference (%)
H3
H5 / β / H7
H10 / H11
H12
0.5 min
2 min
10 min
30 min

### Figure S11

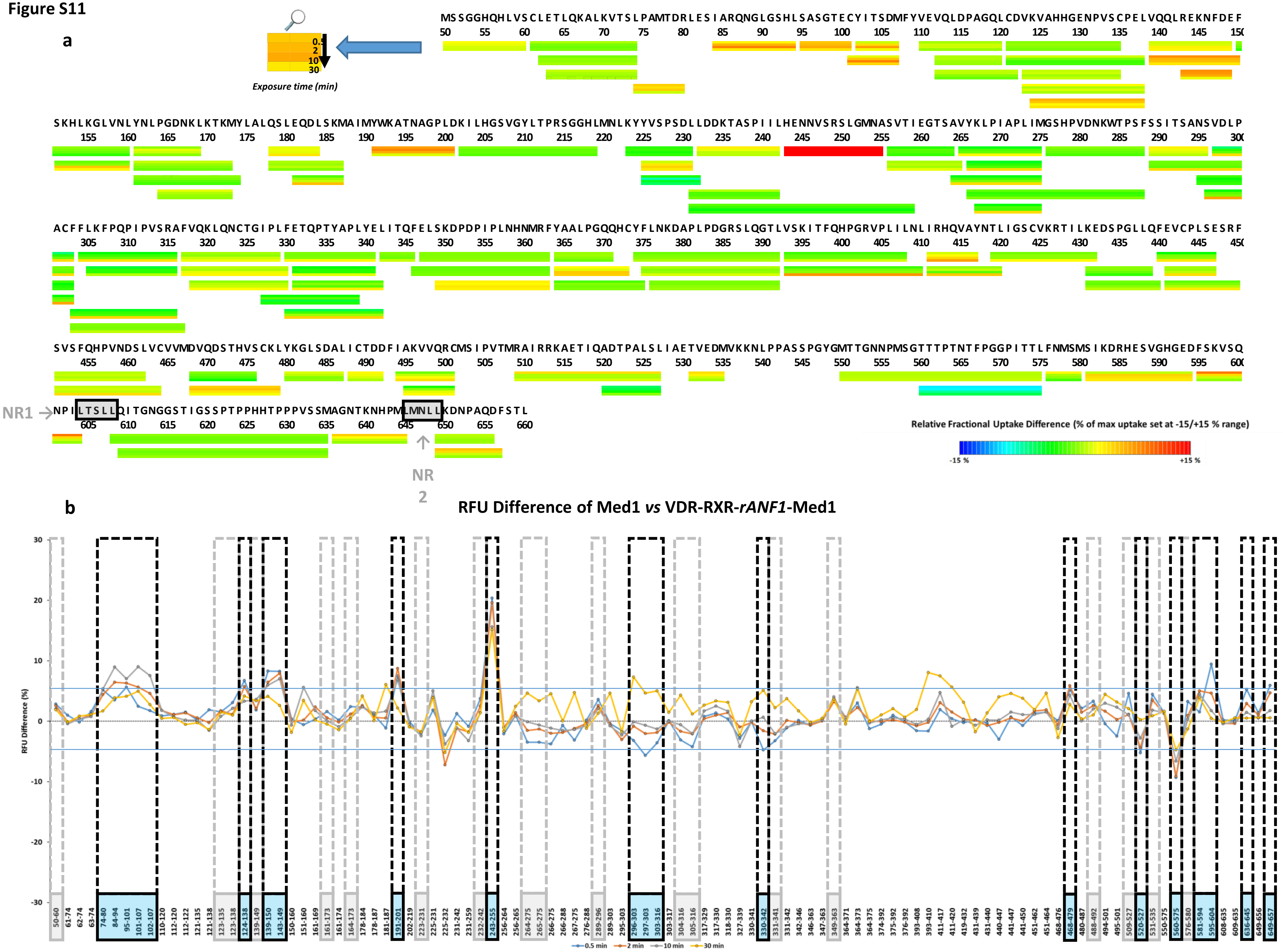

a
Exposure time (min)
0.5
2
10
30
NR1
NR 2
Relative Fractional Uptake Difference (% of max uptake set at -15/+15 % range)
-15 %
+15 %
b
RFU Difference of Med1 vs VDR-RXR-rANF1-Med1
RFU Difference (%)
0.5 min
2 min
10 min
30 min

**Figure S12**

**a**

| Crosslinked proteins | Linked amino acids | Observed crosslink with | | Cα-Cα distance (Å) calculated with PyMOL |
|---|---|---|---|---|
| | | DSBU | C2-arm DSBU | |
| VDR - RXR | K117 – K212 | √ | √ | 10.9 |
| | K109 – K164 | √ | √ | 7.0 |
| | K117 – K164 | √ | √ | 14.8 |
| | K117 – K404 | √ | - | 22.2 |
| | K123 – K212 | √ | √ | 13.5 |
| | K109 – K212 | √ | - | 14.3 |
| | K123 – K164 | √ | - | 21.5 |
| | K55 – K174 | √ | - | 26.0 |
| | K111 – 212 | √ | - | 16.3 |
| | K111 – K164 | √ | √ | 11.2 |
| RXR - RXR | K180 – K193 | √ | - | Unassigned |
| | K212 – K406 | √ | √ | |
| | K200 – K145 | √ | √ | |
| | K132 – K145 | √ | √ | |
| | K180 – K132 | √ | √ | |
| | K406 – K363 | √ | √ | |
| | K212 – K164 | √ | √ | |
| | K164 – K174 | - | √ | |
| | K180 – K164 | - | √ | |
| VDR - VDR | K109 – K117 | √ | √ | Unassigned |
| | K109 – K123 | √ | √ | |
| | K111 – K382 | √ | - | |
| | K111 – K55 | √ | - | |
| | K70 – K111 | √ | - | |
| | K53 – K111 | √ | - | |
| | S168 – K302 | √ | √ | |
| | K53 – K123 | √ | - | |
| | S172 – K302 | - | √ | |
| | K399 – S304 | - | √ | |
| | S165 – K302 | √ | √ | |
| MED 1 – MED 1 | K144 – K69 | - | √ | N.A. |
| | K169 – K393 | √ | - | |
| | K425 – K234 | √ | - | |
| | K495 – K513 | √ | - | |
| | S228 – K425 | - | √ | |
| | K169 – K154 | - | √ | |
| MED 1 – RXR | T236 – K321 | √ | - | N.A. |

**b**

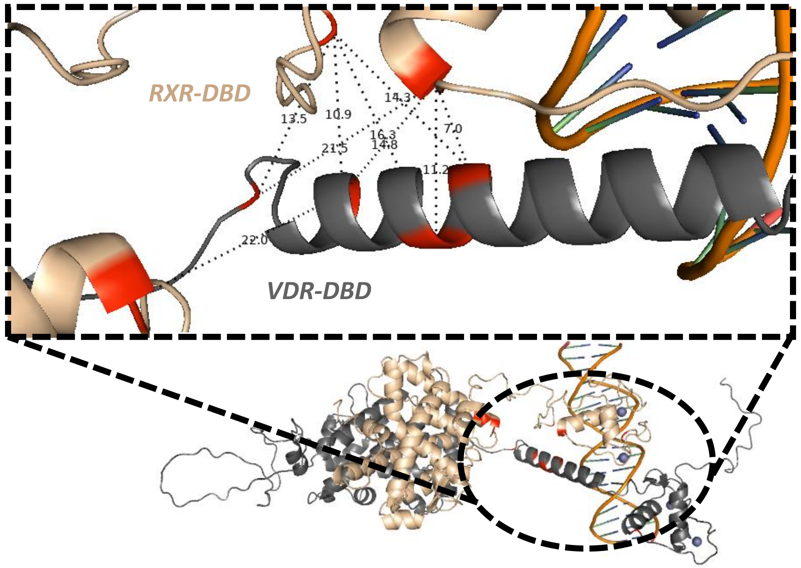
